# Genome-Wide Detection of Imprinted Differentially Methylated Regions Using Nanopore Sequencing

**DOI:** 10.1101/2021.07.17.452734

**Authors:** Vahid Akbari, Jean-Michel Garant, Kieran O’Neill, Pawan Pandoh, Richard Moore, Marco A. Marra, Martin Hirst, Steven J.M. Jones

**Affiliations:** Canada’s Michael Smith Genome Sciences Centre, British Columbia Cancer Agency, Vancouver, British Columbia, Canada; Department of Medical Genetics, University of British Columbia, Vancouver, British Columbia, Canada; Department of Microbiology and Immunology, Michael Smith Laboratories, University of British Columbia, Vancouver, British Columbia, Canada

## Abstract

Imprinting is a critical part of normal embryonic development in mammals, controlled by defined parent-of-origin (PofO) differentially methylated regions (DMRs) known as imprinting control regions. As we and others have shown, direct nanopore sequencing of DNA provides a mean to detect allelic methylation and to overcome the drawbacks of methylation array and short-read technologies. Here we leverage publicly-available nanopore sequence data for 12 standard B-lymphocyte cell lines to present the first genome-wide mapping of imprinted intervals in humans using this technology. We were able to phase 95% of the human methylome and detect 94% of the well-characterized imprinted DMRs. In addition, we found 28 novel imprinted DMRs (12 germline and 16 somatic), which we confirmed using whole-genome bisulfite sequencing (WGBS) data. Analysis of WGBS data in mus musculus, rhesus macaque, and chimpanzee suggested that 12 of these are conserved. We also detected subtle parental methylation bias spanning several kilobases at seven known imprinted clusters. These results expand the current state of knowledge of imprinting, with potential applications in the clinic. We have also demonstrated that nanopore long reads, can reveal imprinting using only parent-offspring trios, as opposed to the large multi - generational pedigrees that have previously been required.

## Introduction

The addition of a methyl group to the 5-carbon of cytidine is the most prevalent and stable epigenetic modification of human DNA (Laurent et al., 2010). DNA methylation is involved in gene regulation and influences a vast array of biological mechanisms, including embryonic development and cell fate, genome imprinting, X-chromosome inactivation, and transposon silencing (Moore et al., 2013; Smith and Meissner, 2013). In mammals, there are two copies or alleles of a gene, one inherited from each parent. Most gene transcripts are expressed from both alleles. However, there is a subset of genes which are expressed from a single allele either randomly such as in X-inactivation, or based upon PofO. The latter is known as imprinting (Chess, 2013; Khamlichi and Feil, 2018).

In imprinting, mono-allelic expression of a gene or cluster of genes is controlled by a *cis-acting* imprinting control region (ICR) (Bartolomei and Ferguson-Smith, 2011). The main mechanism by which this occurs is PofO-defined differential methylation at ICRs, also known as imprinted differentially methylated regions (DMRs) (Bartolomei and Ferguson-Smith, 2011; Maupetit-Méhouas et al., 2016). ICRs are classified as germline (or primary) or somatic (or secondary), hereinafter referred to as gDMR and sDMR. Germline ICRs are established during the first wave of methylation reprogramming at germ cell development and escape the second methylation reprogramming after fertilization (Zink et al., 2018). Secondary or somatic ICRs are established *de-novo* after fertilization during somatic development, usually under the control of a nearby primary ICR (Zink et al., 2018). Imprinted clusters of genes may span up to ∼4 Mb, by acting through a CCCTC-binding factor (CTCF) binding site or by allelic expression of a long non-coding RNA (Bartolomei and Ferguson-Smith, 2011; da Rocha and Gendrel, 2019). By contrast, individually-imprinted genes are typically regulated by PofO-derived differential methylation at the gene promoter (Bartolomei and Ferguson-Smith, 2011).

Imprinting is implicated in various genetic disorders, either from aberrations in imprinting itself, or from deleterious variants affecting the expressed allele at an ICR. Loss of imprinting is also widely observed in human cancers (Goovaerts et al., 2018; Jelinic and Shaw, 2007; Tomizawa and Sasaki, 2012). Thus, accurate mapping and characterization of ICRs in humans is key to the treatment and actionability of genetic disorders, and to personalized oncogenomonics.

To detect ICRs, accurate assignment of methylation data to paternal and maternal alleles is required. Achieving this with traditional bisulfite sequencing or arrays is challenging. Several studies have used samples with large karyotypic abnormalities, such as uniparental disomies (UPDs), teratomas, and hydatidiform moles, to infer regions of imprinting [14–16]. This approach relies on rare structural variants, but also on the assumption that both normal methylation and the imprinted state remain intact in spite of substantial genomic aberrations. A much larger study by Zink *et al*. leveraged a genotyped, multi-generation pedigree spanning nearly half the population of Iceland (n=150,000), in combination with whole genome oxidative bisulfite sequencing (oxBS-Seq), to phase methylation and infer parent-of-origin (Zink et al., 2018). However, despite being able to phase nearly every SNP in that cohort, they were only able to phase 84% of CpG methylation (CpGs on chromosomes 1-22) in over 200 samples due to the short length of reads. Further, that study was based on a single, genetically-isolated population, which may not be representative of the wider human population. A comprehensive mapping of ICRs using a technology more suited to phasing reads, based on individuals more representative of the human population, could greatly advance our understanding of imprinting, with direct benefits for human health.

We have previously shown that nanopore sequencing can detect allelic methylation in a single sample and accurately determine PofO using only trio data. We also developed the software NanoMethPhase for this purpose (Akbari et al., 2021). Here, we applied NanoMethPhase to public nanopore data from a diverse set of 12 lymphoblastoid cell lines (LCLs) from the 1,000 Genomes Project (1KGP) and Genome in a Bottle (GIAB) to investigate genome-wide allele-specific methylation (ASM) and detect novel DMRs (Figure 1A) (Auton et al., 2015; De Coster et al., 2019; Jain et al., 2018; Shafin et al., 2020; Zook et al., 2019, 2016). Using trio data from 1KGP for these cell lines we phased nanopore long reads to their PofO and inferred allelic methylation (Akbari et al., 2021; Auton et al., 2015). Nanopore was able to detect haplotype and methylation status for 26. 5 million autosomal CpGs (Chromosomes 1-22), which represents 95% of the autosomal CpGs in the GRCh38 (Kent et al., 2002). We further used public whole-genome bisulfite sequencing (WGBS) data to confirm the presence of the detected DMRs in other tissues and to class the novel DMRs as being germline or somatic. We captured 94% of the well-characterized DMRs (Those reported by at least two studies) and detected 28 novel DMRs (12 germline and 16 somatic). We determined that 43% of these novel DMRs show evidence of conservation in rhesus macaque and chimpanzee. Collectively, our results extend the set of known imprinted intervals in human and demonstrate a major contribution in our ability to characterize imprinting by ASM, brought about by the capabilities of nanopore sequencing.

**Figure 1:**
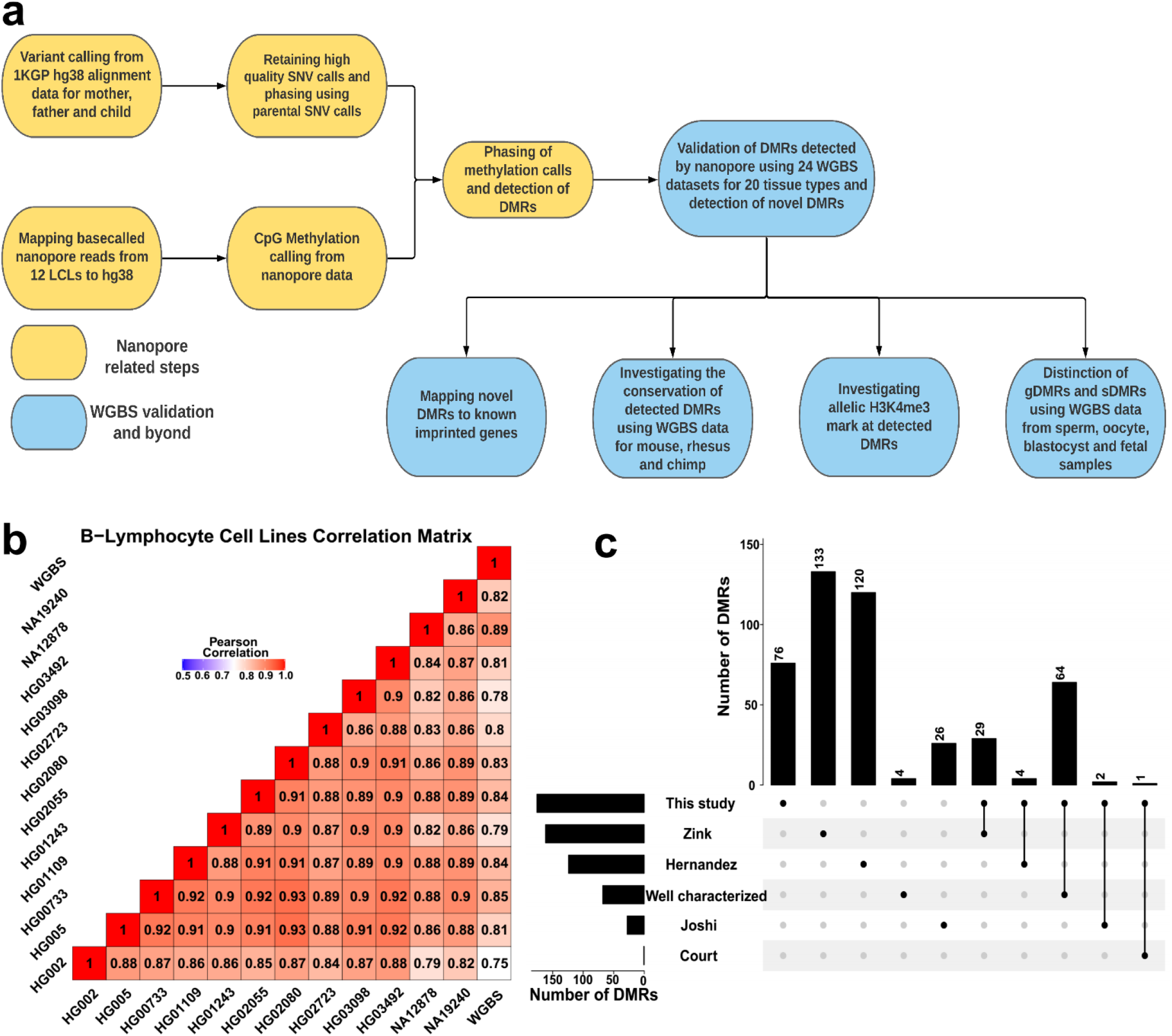
Detection of allelic methylation using nanopore sequencing. a) The flowchart of the study representing all the analysis steps. b) Pearson correlation matrix of the nanopore CpG methylation frequencies for the 12 LCLs and NA12878 whole-genome bisulfite sequencing (WGBS) from ENCODE (ENCFF835NTC). c) Upset plot of the number of DMRs detected in our study and previous studies and their overlaps.

## Results

### Assessing the Effectiveness of Nanopore Methylation Calling and Detection of Known Imprinted DMRs

We performed correlation analysis among cell lines and NA12878 nanopore-called methylation with WGBS data (ENCFF835NTC) to confirm the reliability of methylation calling (Figure 1B). We observed high correlation across cell lines (*r* = 0.75-0.93), as expected due to their being the same cell type. NA12878 nanopore-called methylation also showed the highest correlation (*r = 0*.*89*) with NA12878 WGBS (Figure 1B), as expected. Additionally, to examine performance on detection of known DMRs, we gathered the list of reported DMRs from previous studies (Court et al., 2014; Hernandez Mora et al., 2018; Joshi et al., 2016; Zink et al., 2018). This included 383 imprinted intervals, of which 68 we assigned as “well-characterized” because they were reported by at least two genome-wide mapping studies or were previously known to be imprinted (Supplementary file1). Subsequently, we haplotyped the methylome in each cell line, performed differential methylation analysis (DMA) between alleles across cell lines. 95% (26.5M) of human autosomal CpGs could be assigned to a haplotype. We detected 172 allelic DMRs (p - value < 0.001, |methylation difference| > 0.25, and detected in at least 4 cell lines in each haplotype). See supplementary file 2 for more details. Of the 172 detected DMRs, 96 (56%) overlapped with at least one previously reported, while the remaining 76 (44%) were novel. Of the well-characterized DMRs, 64/68 (94%) were detected in our study (Figure 1C, supplementary file2). All DMRs which overlapped with previously-reported DMRs displayed consistent PofO with those studies.

We similarly examined the power of nanopore sequencing to detect allelic DMRs within a single sample, by comparing to previous studies (Court et al., 2014; Hernandez Mora et al., 2018; Joshi et al., 2016; Zink et al., 2018). On average, 88% (M ± SD = 24.5M ± 1.7M) of the human methylome could be assigned to a parental haplotype in each LCL. Of the well-characterized DMRs, ∼71% (M ± SD = 48 ± 4.8) could be detected in a single LCL. An additional 32 DMRs (SD = 9.7) reported by only one previous study were detected in each sample (Supplementary figure S1).

### Confirmation of Novel Imprinted DMRs

As noted above, we detected 76 allelic DMRs that did not overlap with previously-reported ICRs (Court et al., 2014; Hernandez Mora et al., 2018; Joshi et al., 2016; Zink et al., 2018). In order to determine their validity as novel DMRs, we used 24 WGBS datasets from 20 tissue samples within the Roadmap Epigenomics Project (See materials and methods) (Bernstein et al., 2010). We first examined the 96 allelic DMRs which overlapped with the reported DMRs. 79 out of 96 DMRs that overlapped with reported regions showed adjusted p-value (FDR) < 5 × 10^−6^ and log fold change > 0.15, while only 5, 6, 7, and 8 intervals were detected as significant in the control intervals including 200 randomly selected 1kb bins, CpG islands, 2kb, and 3kb bins, respectively (Figure 2A, Supplementary file 3). Applying this approach to the 76 not previously reported DMRs, the WGBS data supported 28 significant DMRs (Figure 2A and 2B, Supplementary file 3). In agreement with previous studies reporting higher number of maternally methylated intervals, 10 of the 28 novel DMRs were paternally methylated and 18 were maternally methylated (Court et al., 2014; Hernandez Mora et al., 2018; Joshi et al., 2016). Overall, 107 out of 172 DMRs were validated in tissue WGBS data from which 28 were novel and 79 were reported by the previous studies (Figure 2C, Supplementary file 2) (Court et al., 2014; Hernandez Mora et al., 2018; Joshi et al., 2016; Zink et al., 2018).

**Figure 2:**
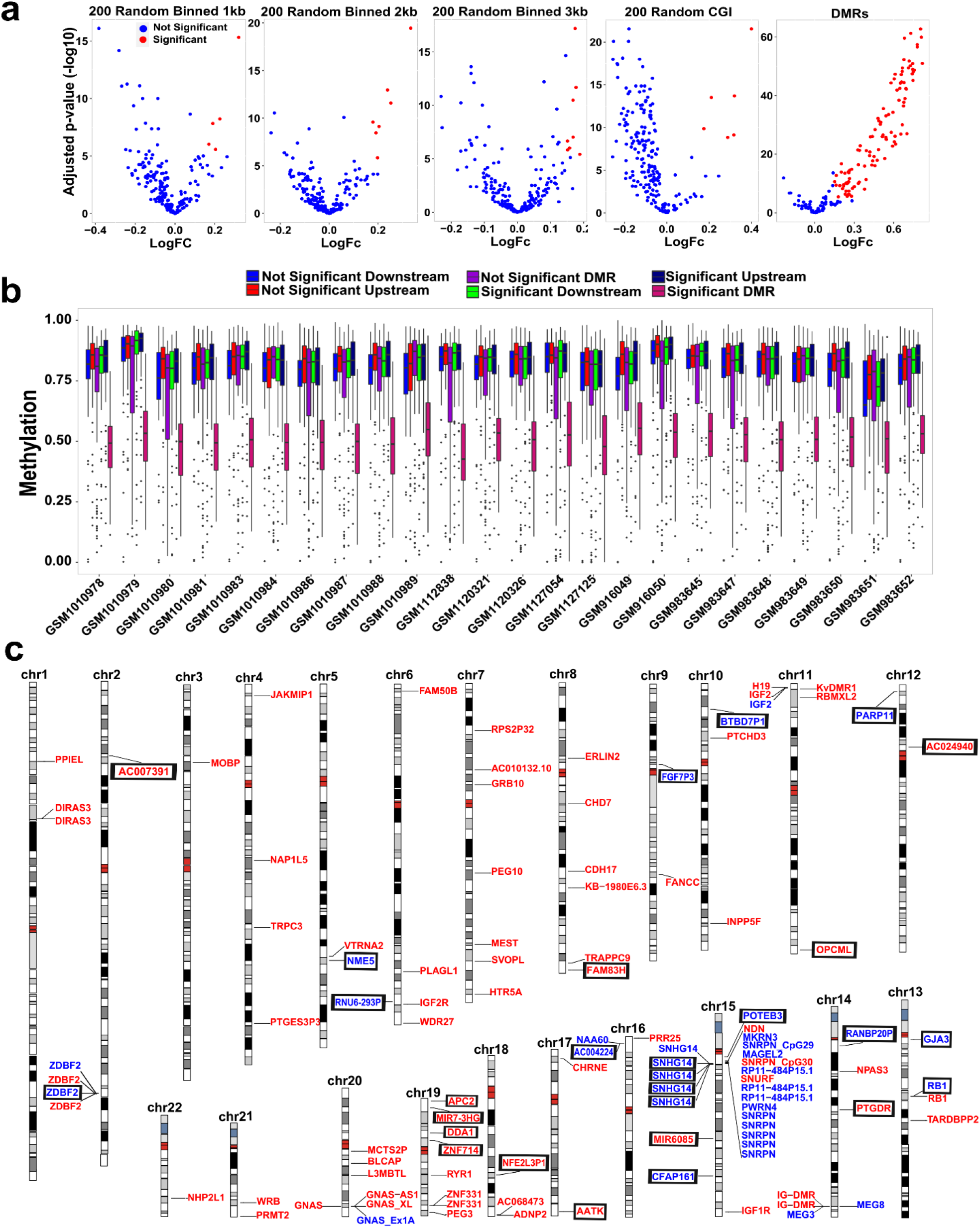
Validation of nanopore-detected DMRs using WGBS data. a) Volcano plots representing limma results for partial methylation (30% -70%) analysis of nanopore-detected DMRs in 24 WGBS datasets from 20 tissue types. 200 randomly selected CpG islands, 1kb, 2kb, and 3kb intervals are examined as controls. Red dots adjusted p-value (FDR) < 5 × 10^−6^ and log fold change > 0.15. b) Box plot showing partial methylation at significant DMRs while not significant DMRs and adjacent regions (down- and upstream to significant and not significant DMRs) are not partially methylated. c) Idiogram of the 107 DMRs which validated by WGBS. On the left on each chromosome are paternally methylated DMRs and on the right are maternally methylated DMRs. Red color represents gDMRs and blue represents sDMRs. Novel DMRs are boxed and named based on their nearest gene (Ensembl Gene 103 GRCh38.p13).

We also sought to examine the significance of the other 283 previously reported imprinted regions which did not overlap with our detected DMRs. We examined these 283 DMRs in WGBS data and only 139/283 DMRs (49%) were significant (adjusted p-value (FDR) < 5 × 10^−6^ and log fold change > 0.15. Supplementary file 4). We also mapped these 283 intervals to the DMRs detected in each LCL sample. 81/283 (27%) of them were detected in at least one sample with consistent reported PofO, of which 41 were in common with WGBS analysis (Supplementary file 5).

### Determination of Germline vs Somatic Status of Novel Imprinted DMRs

We performed DMA between oocyte and sperm and overlapped detected DMRs to the 28 novel DMRs. 12 of the novel DMRs overlapped with DMRs from oocyte versus sperm (p-value < 0.001, |methylation difference| > 0.25, and more than 40% methylation in oocyte and less than 20% in sperm and vice versa) from which 11 were maternally methylated and 1 was paternally methylated (Figure 3). We then examined the methylation of somatic and germline DMRs in early human embryonic cells and fetal tissues to investigate whether the imprinting of the 12 candidate gDMRs survived the second round of de- and remethylation and if the other 16 novel sDMRs were established during development. We used blastocyst WGBS data from early cleavage-stage embryos and fetal tissue (Bernstein et al., 2010; Okae et al., 2014). All novel candidate gDMRs showed partial methylation in the blastocyst indicating the gDMRs escaped de-methylation after fertilization (Figure 3). All novel gDMRs and sDMRs displayed partial methylated in fetal tissues indicating survival of gDMRs during somatic development and establishment of sDMRs. Overall, 12 of the novel DMRs detected to be germline while 16 detected as sDMRs (Figure 2C and Figure 3, Supplementary file 6).

**Figure 3:**
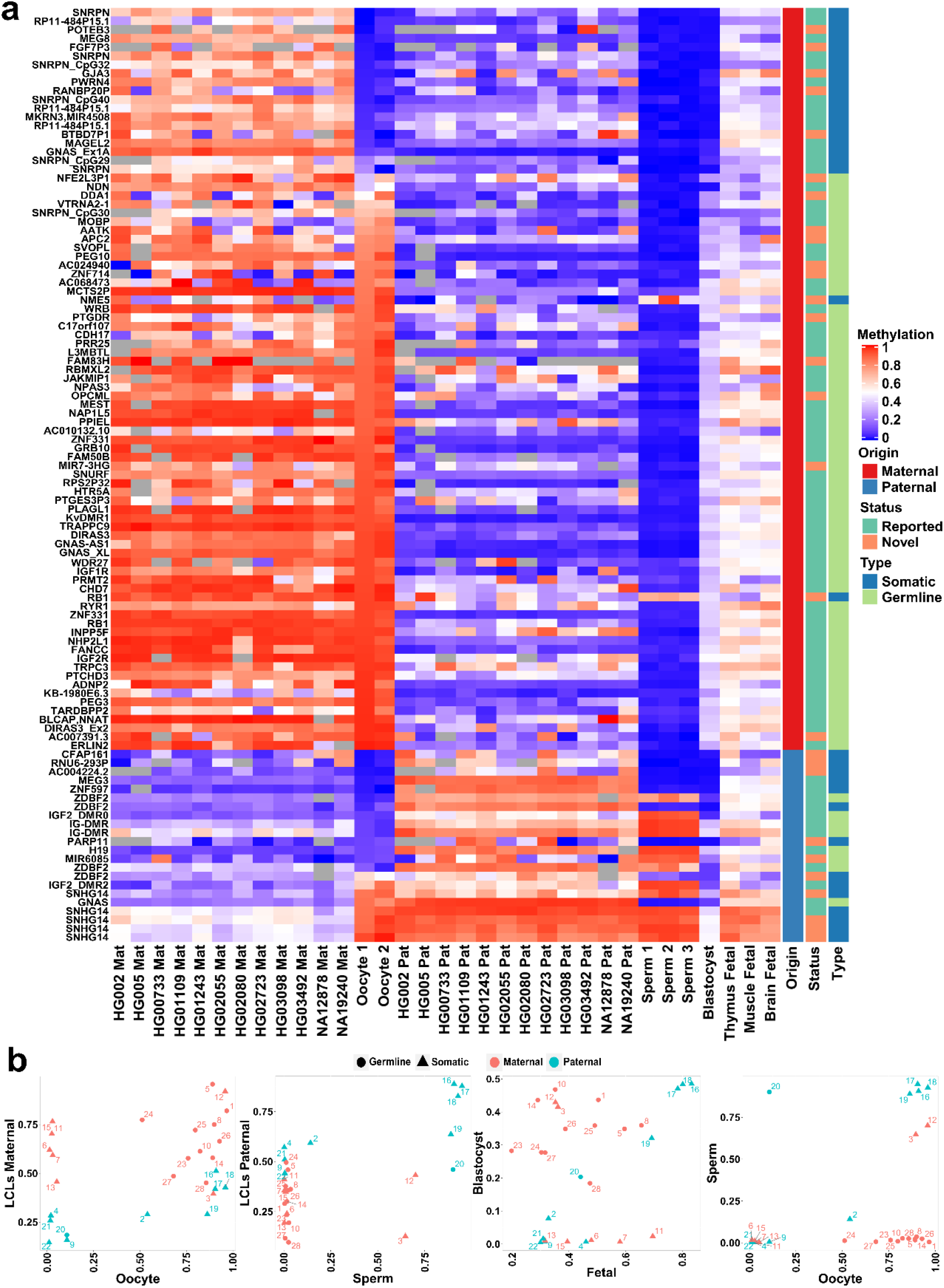
Detection of novel gDMRs and sDMRs. a) Heatmap displaying average methylation of the 107 nan opore-detected DMRs which validated by WGBS. Average allelic methylation at each DMR for LCL samples are represented. Average methylation of each DMR for gametes and early developmental WGBS samples are displayed. Novel DMRs are named based on their nearest gene (Ensembl Gene 103 GRCh38.p13). b) Dot plots representing the methylation of novel gDMRs and sDMRs in each sample in respect to other sample. Maternally methylated g DMRs display high methylation in oocyte and very low or no methylation in sperm and are partially methylated in blastocyst and fetal samples. Paternally methylated gDMRs display high methylation in sperm and very low or no methylation in oocyte and are partially methylated in blastocyst and fetal samples. Somatic DMRs do not display relevant methylation in sperm or oocyte and are methylated or unmethylated in both sperm and oocyte while they display parental methylation bias in LCLs and partial methylation in fetal samples. For ease in visualization DMR ID s are shown (Supplementary file 6).

### Allelic Histone Methylation of H3K4 is Enriched at Germline DMRs

The H3K4me3 histone mark is protective to DNA methylation. At ICRs, the unmethylated allele is usually enriched for this histone modification (Court et al., 2014; John and Lefebvre, 2011). We used H3K4me3 chromatin immunoprecipitation sequencing (ChIP-seq) data for 7 LCLs and their heterozygous single-nucleotide variant (SNV) calls from 1KGP. 81/107 of the detected DMRs could be examined (See material and methods). Of these, 42 reported and 9 novel DMRs showed a significant allelic count in ChIP-seq data (Fisher’s combined p-value binomial < 0.01) (Supplementary files 6 and 7). Among the 7 LCLs with ChIP-seq data, only NA12878 and NA19240 were among LCLs with nanopore data and a phased methylome. Therefore, we examined if the allelic H3K4me3 and methylation are in opposite alleles in these cell lines. 23 reported and 5 novel DMRs were significant for allelic H3K4me3 in NA12878 and/or NA19240. 21 reported and 4 novel DMRs showed opposite allelic bias state between H3K4me3 and methylation (Supplementary file 7).

Allelic H3K4me3 mostly overlapped with gDMRs. Overall, 75% of assessable gDMRs and 39% of sDMRs were significant for allelic H3K4me3. This is consistent with previous studies demonstrating the protective role of H3K4me3 against DNA methylation, specifically at germline ICRs in the second round of re-methylation during implantation and somatic development (Chen and Zhang, 2020; Hanna and Kelsey, 2014).

### Conservation of Detected Imprinted DMRs across Mammals

To investigate the conservation of detected DMRs and determine if any of the novel DMRs are conserved in mammals we used WGBS data from house mouse (Mus musculus), rhesus macaque (Macaca mulatta), and chimpanzee (Pan troglodytes) (Hon et al., 2013; Jeong et al., 2021; Tung et al., 2012). We examined whether any of the orthologous regions in these mammals display significant partial methylation (Materials and Methods). Of the 107 DMRs detected by nanopore and validated in WGBS data, 71, 105 and all 107 had orthologs in mouse, rhesus and chimp, respectively. Orthologs of the 77/107 detected DMRs showed significant partial methylated in at least one of the three mammals (Figure 4A, Supplementary file 6 and 8). Of these, 65 were reported DMRs (56 well-characterized) and 12 novel DMRs. We detected 24 significant orthologous DMRs in mouse. 18 of these were reported to be imprinted by previous studies in mouse (Gigante et al., 2019; Xie et al., 2012). All significant DMRs in mouse, except one, were also significant in rhesus and/or chimp suggesting their existence in their common ancestor. These DMRs mapped to well-known imprinted clusters including *KCNQ1, H19, GNAS*, SNURF/*SNRPN, PLAGL, SGCE, BLCAP, PEG3, PEG10, PEG13, GRB10, BLCAP, NAP1L5, INPP5F*, and *MEG3* where their allelic PofO expression has already been reported in mouse and other mammals (“Geneimprint,” 2021; Morison et al., 2001).

**Figure 4:**
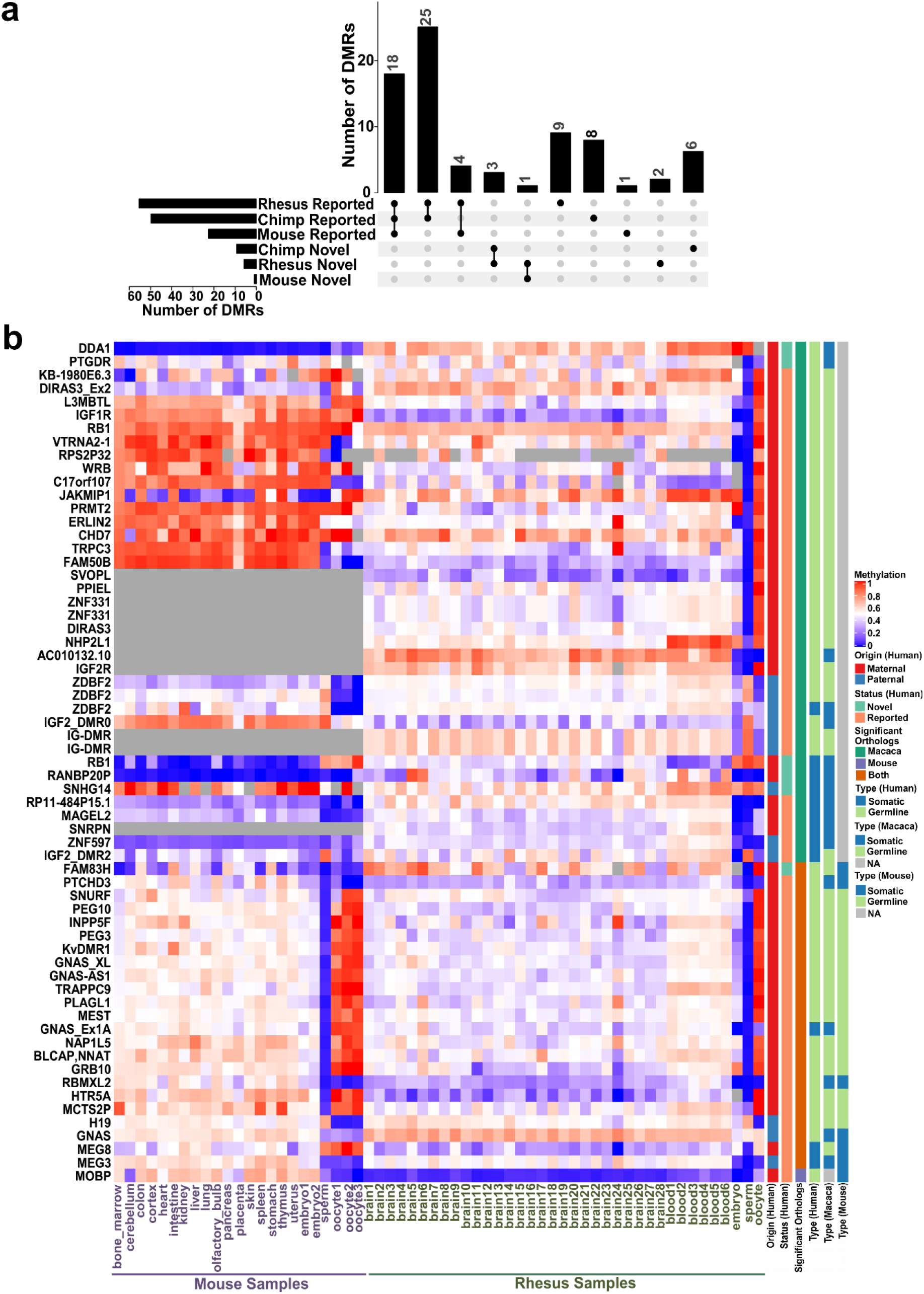
Conservation of detected DMRs. a) Upset plot representing the number of reported and novel DMRs with evidence of conservation (partial methylation) in each of the mammals. b) Heatmap representing human DMRs (DMR names on the left of the heatmap) and average methylation of their orthologous intervals in mouse and macaca in different tissues and also in sperm, oocyte, and embryonic samples.

Sperm, oocyte and embryo WGBS data for mouse and rhesus were used to investigate if the DMRs that detected as germline or somatic in human are germline or somatic in these mammals and vice versa (Dahlet et al., 2020; Gao et al., 2017; Jung et al., 2017; Saenz-de-Juano et al., 2019). 62 of the human DMRs had significant orthologs in rhesus. Of these, 51 were germline and 11 were somatic in human and in rhesus 45 were germline and 17 were somatic (More than 40% methylation in oocyte and less than 20% in sperm and vice versa with |methylation difference| > 0.25. Figure 4B). 24 human DMRs had significant orthologs in mouse. Of these, 21 were germline and 3 were somatic in human and in mouse 17 were germline and 7 were somatic (Figure 4B). Nine gDMRs in human were somatic in rhesus and/or mouse and three gDMRs from mouse or rhesus were somatic in human. This is consistent with previous studies indicating imprinting is largely conserved in mammals while ICR identity at the germline stage is not completely conserved (Cheong et al., 2015).

### Novel DMRs within Known Imprinted Gene Domains and Contiguous Blocks of Parental Methylation Bias

We gathered the list of 259 imprinted genes from previous studies (Supplementary file 9) (Babak et al., 2015; Baran et al., 2015; “Geneimprint,” 2021; Jadhav et al., 2019; Morison et al., 2001; Zink et al., 2018). 14 novel DMRs (6 germline and 8 somatic) mapped close (<1.03Mb) to imprinted genes (Supplementary file 6 and 10).

Of the 8 sDMRs close imprinted genes, only one mapped to a CpG island, and that was a small (<300bp) CpG island ∼13 Kb downstream of the maternally expressed *NAA60* gene (Supplementary figure S2). Four novel sDMRs (All paternally methylated) mapped in the Prader-Willi syndrome and Angelman syndrome (PWS/AS) cluster. Previous studies reported continuous subtle paternal methylation bias at the PWS/AS cluster (Hernandez Mora et al., 2018; Joshi et al., 2016; Zink et al., 2018). Consistent with previous studies, the four novel sDMRs at this cluster were large (>5Kb) and seemed to constitute near-continuous paternal methylation spanning a ∼200kb region. This included the *SNORD116* cluster genes and several other genes such as *PWAR1* and *6, PWARSN* and *IPW* (Supplementary figure S3). This paternally methylated somatic block is downstream of the maternally methylated germline *SNURF/SNRPN* ICR, which is associated with PWS and shows evidence of conservation in chimp, rhesus, and mouse. Moreover, the allele-specific expression (ASE) track from Zink *et al*. displayed strong paternal expression across this ∼200kb region (Zink et al., 2018). Another three novel sDMRs mapped close *RB1/LPAR6, IGF2R* (Supplementary figures S4 and S5) and *GPR1-AS/ZDBF2*. The novel sDMR at *GPR1-AS/ZDBF2* were close to 2 known paternal gDMRs. Moreover, LCLs PofO methylation track at the *ZDBF2* gene body showed continuous subtle paternal bias. Together, these suggest a ∼65kb paternally methylated block interrupted by unmethylated CpG island at *ZDBF2* promoter (Supplementary figure S6). In addition to blocks with novel DMRs, we sought to detect continuous block of parental methylation bias at other regions. We detected 5 other contiguous blocks of imprinting at *ZNF331, KCNQ1OT1, GNAS, L3MBTL1* and *ZNF597/NAA60*, ranging from 35-58Kb in size (Supplementary figures S7-11).

All the six gDMRs within imprinted gene domains were maternally methylated and they all mapped to CpG islands except a DMR mapped in the *AC024940*.*1* (*OVOS2*) (Supplementary figures S12-16, Figure 5). Five of them mapped to known imprinted genes without previously reported DMR or a DMR with a much greater distance from the gene compared to our DMRs including *AC024940*.*1, ZNF714, DDA1, ADAMTSL5*, and *NAPRT* (Court et al., 2014; Hernandez Mora et al., 2018; Joshi et al., 2016; Zink et al., 2018). A novel gDMR mapped to the promoter of *ZNF714* which is reported to be paternally expressed (Jadhav et al., 2019; Zink et al., 2018). Thus suggesting this DMR could be the potential ICR and directly suppress maternal allele by blocking its promoter (Figure 5). *AC024940*.*1* reported to be paternally expressed (Zink et al., 2018). A novel germline maternal DMR mapped near the end of the *AC024940*.*1* gene (encompassing the intron 38 to the start of exon 40) adjacent to a CTCF binding site (Supplementary figure S12). *DDA1* and *ADAMTSL5* have been previously reported to be maternally expressed and *NAPRT* has an isoform dependent expression origin (Babak et al., 2015; Zink et al., 2018). A gDMR mapped to the end and downstream of *DDA1* gene (Supplementary figure S13). For *ADAMTSL5* and *NAPRT*, gDMRs mapped close to these genes (<150Kb) (Supplementary figures S14 and S15).

**Figure 5:**
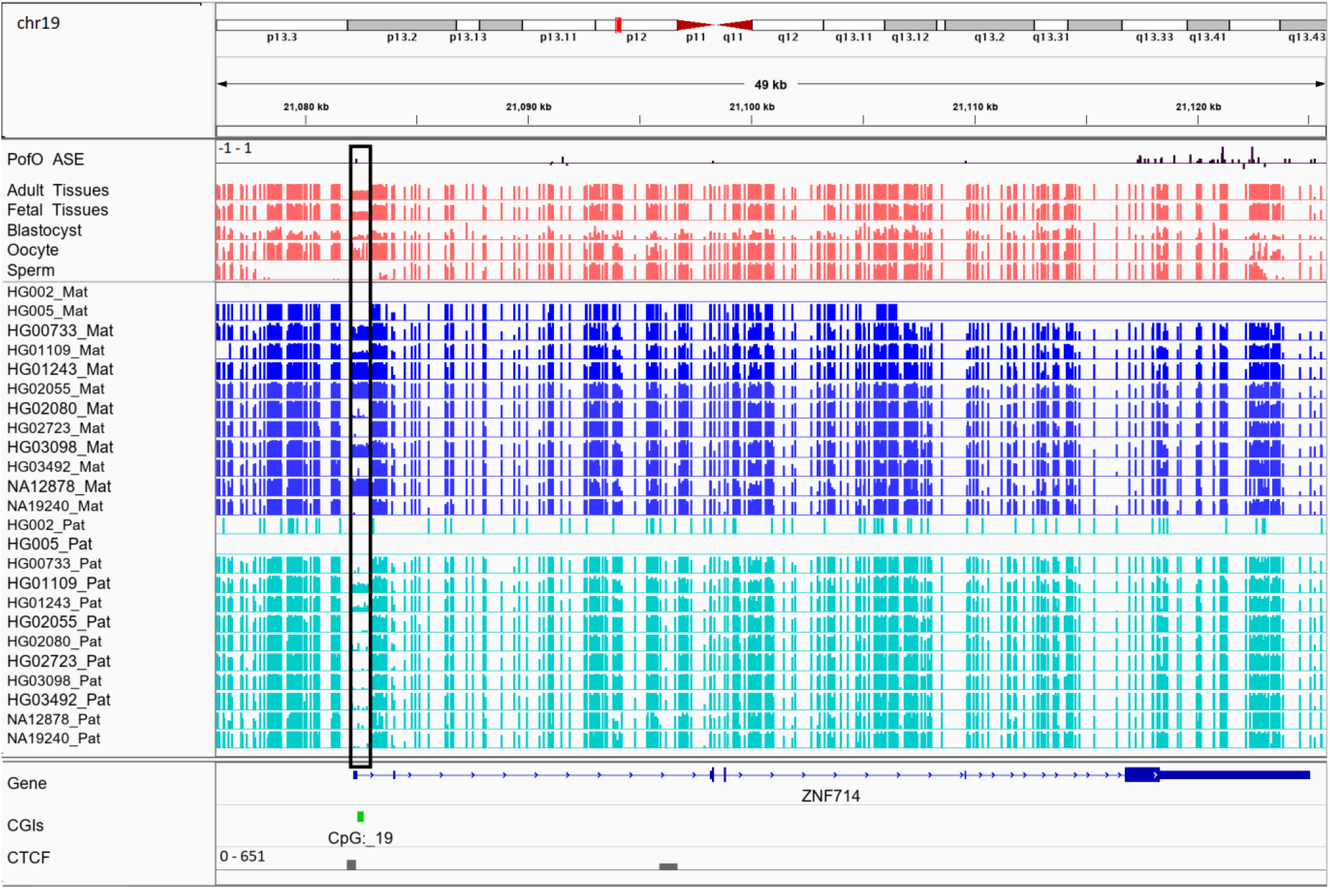
IGV screenshot of the novel maternally methylated germline DMR at the promoter of the paternally expressed ZNF714 gene. Black box region represents the DMR. PofO_ASE represents allele specific expression track from Zink *et al*. without any filtering for *P value* and positive vertical bars (upward) represents more paternal expression and negative bars (downward) maternal expression. The range for all methylation tracks is 0 -1. Adult tissues track represents the average methylation of 24 WGBS data from 20 tissue samples and fetal tissues track is the average methylation of 3 fetal WGBS tissue samples. These WGBS samples obtained from Roadmap Epigenomics Project. Blastocyst, oocyte and sperm tracks are the methylation from one blastocyst, and average methylation from two oocytes, and three sperm samples from Okae et al., 2014.

## Discussion

Here we described the first genome-wide map of human allele-specific methylation using nanopore sequencing. Leveraging long reads and parental SNVs allowed us to phase methylation for ∼26.5 million autosomal CpGs representing 95% of the CpGs in the human autosomal genome (GRCh38) across 12 LCLs (De Coster et al., 2019; Kent et al., 2002; Shafin et al., 2020; Zook et al., 2016). This represents a much higher resolution than previous studies aimed to capture allelic methylation (Court et al., 2014; Hernandez Mora et al., 2018; Joshi et al., 2016; Zink et al., 2018). For example, Zink *et al*. determined the PofO of almost all genotypes using nearly half the population of Iceland (n=150,000) and used over 200 whole-genome OxBS-seq samples to detect imprinting (Zink et al., 2018). They could define PofO methylation for ∼23.5 million autosomal CpGs (84%). We noticed three of our novel DMRs did not have any CpG representation from Zink *et al*. Moreover, in a further three other novel DMRs, fewer than 60% of the CpGs were captured in Zink *et al*. (Supplementary file 6). EPIC methylation arrays detect over 850k CpGs and covers almost all CpGs detectable by 450k and 27k methylation arrays. Seven of our novel DMRs did not have any CpGs covered by the EPIC array and 9 other novel DMRs had only 1 or 2 probes on this platform (Supplementary file 6). This highlights the breadth of nanopore sequencing for the purposes of ICR calling.

Even though we detected methylation for all the CpGs in the human genome (GRCh38), we were not able to phase 5% of the human methylome (Kent et al., 2002). To phase nanopore reads, we used SNVs detected from short-reads data in the 1KGP (Auton et al., 2015). Short-reads are challenging to map to complex repetitive regions which results in lack of SNVs and subsequent inability to phase reads in these regions. 75% of the unphased CpGs mapped to the ENCODE blacklist (Amemiya et al., 2019). We previously demonstrated that using SNVs detected from nanopore to phase reads results in reliable methylation phasing and detection of a few more reported DMRs (Akbari et al., 2021). Improvement in basecalling and variant calling from nanopore reads could enable the phasing of a complete genome-wide methylome using nanopore detected SNVs.

Using nanopore sequencing we could capture 94% of the well-characterized DMRs and 35 of the DMRs reported by only one study (Court et al., 2014; Hernandez Mora et al., 2018; Joshi et al., 201 6; Zink et al., 2018). However, we were unable to detect a further 283 DMRs, mostly reported by one previous study (Court et al., 2014; Hernandez Mora et al., 2018; Joshi et al., 2016; Zink et al., 2018). In further analyses, 180 of these DMRs were detected in at least one nanopore-sequenced LCL sample and/or validated in the Roadmap multi-tissue WGBS data we used (Supplementary files 4 and 5). We should note that nanopore data comes from a small number of B-lymphocyte cell-line samples, yet considerably diverse in ethnicity. Imprinted DMRs can be tissue-specific and polymorphic across individuals, which may explain this discrepancy (Court et al., 2014; Hernandez Mora et al., 2018; Joshi et al., 2016; Romanelli et al., 2014; Silver et al., 2015; Zink et al., 2018). Characterization of imprinted DMRs across a wider range of tissues and populations represents a clear path forward for the field. The ability of nanopore sequencing to characterize imprinting using only parent-offspring trios represents a relatively low-cost avenue by which this might be achieved.

We detected 107 DMRs using nanopore which were further confirmed in multi-tissue WGBS data. Twelve of these were novel gDMRs and sixteen were novel sDMRs not reported in previous studies (Court et al., 2014; Hernandez Mora et al., 2018; Joshi et al., 2016; Zink et al., 2018). These novel DMRs were supported by several lines of evidence in our analyses. 1) They displayed significant PofO methylation bias in nanopore LCLs. 2) They were significantly partially methylated in WGBS data from 20 human tissues. 3) gDMRs demonstrated escape from the second de-methylation step. 4) They were partially methylated in three fetal tissue samples. 5) 43% of those for which H3K4me3 ChIP-seq data could be phased showed significant allelic H3K4me3. 6) 43% showed evidence of conservation in at least one of the three mammals including chimp, rhesus, and mouse. 7) 71% mapped to at least one regulatory region including CpG island, CTCF binding site and enhancer. These novel DMRs represent a substantial and well-validated expansion of known regions of imprinting, which may aid future research and diagnosis in the fields of genetic medicine and oncology.

Of the 107 DMRs, 20 mapped to the PWS/AS cluster. Previous studies demonstrated two paradigms of imprinting at this cluster, either PofO methylation confined to particular regulatory regions such as CpG islands or subtle paternal bias across this cluster with spikes of maternal methylation (Court et al., 2014; Joshi et al., 2016; Sharp et al., 2010; Zink et al., 2018). Although we did not observe paternal methylation bias across the whole PWS/AS cluster, we did detect a paternal methylation block spanning ∼200Kb, immediately downstream of the known maternally methylated PWS *SNURF/SNRPN* ICR. This block encompasses the *SNORD116* cluster and other adjacent genes with strong paternal expression (Supplementary figure S3). Probes with paternal methylation bias at the *SNORD116* cluster have been reported which span about 95Kb region and paternal deletion of this cluster results in PWS phenotypes (Hernandez Mora et al., 2018; Joshi et al., 2016; Matsubara et al., 2 019). Slight hypomethylation of *SNORD116* cluster in cases with PWS phenotype and hypermethylation in the cases with AS phenotype have been reported (Matsubara et al., 2019). Our analysis extends and more clearly delineates this paternally biased block.

Beyond the PWS/AS cluster, we detected another six blocks of allelic methylation bias (Supplementary figure S6-S11). All of the blocks represented several features in common. 1) They were detected in imprinted genes that appeared in cluster. 2) All of them were accompanied by a strong PofO expression bias from the subtle hypermethylated allele. 3) There was at least one well-characterized and conserved gDMR in each block (except *ZNF597/NAA60* block with a conserved sDMR). 4) The well-characterized DMRs in these blocks displayed significant allelic H3K4me3 (except DMR in *L3MBTL1* block which could not be examined due to the lack of SNV). 5) Well-characterized DMRs in these blocks overlapped to the promoter of genes with subtle PofO methylation bias at the gene body and DMR itself displayed opposite PofO methylation (except for *ZDBF2/GPR1-AS* block that DMR did not mapped to the promoter and had the same PofO with the gene body). This represents a novel facet of imprinting biology. To explain this, we can consider that CpG methylation at gene bodies is positively (but weakly) correlated with gene expression (Ball et al., 2009; Yang et al., 2014). Within these blocks, we saw parental methylation bias at the parentally expressed or active allele. This may suggest that subtle parental methylation is linked to parental ASE. However, ASE is observed in many other imprinted genes whose gene bodies do not show parental methylation bias. One possible explanation could be that the subtle parental methylation bias is used by cells to express important genes (genes which can regulate other genes in the cluster or have regulatory roles) in an imprinted cluster with higher fidelity through its gene body methylation on active allele. For example, at the *KCNQ1OT1* and *GNAS* clusters the methylation blocks overlap *KCNQ1OT1* and *GNAS-AS1* genes both of which encode antisense RNA transcripts that regulate other genes in the imprinted cluster (Chiesa et al., 2012; Turan and Bastepe, 2013). However, further studies are needed to reveal the mechanism producing these contiguous slight parental methylation bias blocks and their functional role.

Orthologous regions of ∼72% of the detected DMRs were demonstrated significant partial methylation in at least one of the chimp, rhesus, and mouse. There were a considerably higher number of orthologous sites and significant orthologous DMRs in chimp and rhesus in agreement with more similarities and less distance to these primates compare to mouse in the human evolution. Orthologs of the 12 novel DMRs were mostly displayed significant partial methylation in rhesus and/or chimp while the other 16 novel DMRs were not significant in any of the examined mammals (Figure 4). This suggests that the novel DMRs (except one which had significant orthologous in mouse) are established after divergence of primates’ common ancestor from mouse and majority of them established after the divergence of human common ancestor from chimp. Court *et al*. detected 14 novel DMRs, at the time of their study, and did not detect any imprinted orthologs of their novel DMRs in mouse (Court et al., 2014). All 14 also overlapped with our detected DMRs and six of them had orthologous regions in mm10 using the UCSC liftover file (Kent et al., 2002). Two of the orthologs displayed partial methylation in mouse, one in *Rian* gene which did not examined in Court *et al*. and the other in *Htr5a* gene which reported not to be conserved in mouse by Court *et al*. (Court et al., 2014). When looking into their analysis, it seems that they examined different orthologous region (Supplementary figure S19). For *Htr5a*, they examined the CpG island (CpG:_102) ∼50 kb away from the gene while we examined the region spanning the first or second exon (two transcripts) of *Htr5a* which was partially methylated while CpG:_102 was also unmethylated in our study.

Using reported imprinted genes, 50% of the novel DMRs mapped close to known imprinted genes (Babak et al., 2015; Baran et al., 2015; “Geneimprint,” 2021; Jadhav et al., 2019; Morison et al., 2001; Zink et al., 2018). Five of our novel gDMRs could be potential ICRs for reported imprinted genes without reported ICR. Specifically, maternal methylation of CpG island overlapping promoter of *ZNF714* as it can directly repress maternal allele and results in the reported paternal expression (Figure 5) (Jadhav et al., 2019; Zink et al., 2018). *ZNF714* is a member of the zinc finger family proteins which have several imprinted genes with developmental roles (Babak et al., 2015; Baran et al., 2015; Camargo et al., 2012; Jadhav et al., 2019; Zink et al., 2018). ZNF714 has been reported to be associated with non-syndromic cleft lip (Camargo et al., 2012). Thus, this new imprinted DMR could be of potential clinical value. In contrast to imprinting which is established in the germline and usually consistent across tissues, allelic expression is only present if the imprinted gene is expressed in the tissue. Moreover, studies have used short read sequencing to detect ASE which is confounded with several limitations (Aird et al., 2011; Steijger et al., 2013). Therefore, a comprehensive ASE analysis using long-read technologies capturing various tissues might explain ASE around the novel DMRs without evidence of a close imprinted gene. Paternal expression bias of *PTCHD3* and maternal expression bias for *FANCC* are detected in Zink *et al*. while they could not detect any associated DMR (Zink et al., 2018). Hernandez *et al*. detected 3 and 1 maternally methylated probes at the promoter of *PTCHD3* and intron one of *FANCC*, respectively, but were not able to examine the parental expression (Hernandez Mora et al., 2018). We also detected two maternally methylated gDMRs overlapping the promoter of *PCTHD3* and intron one of *FANCC* (Supplementary figures S17 and S18). There were no phased CpG for these DMRs in Zink *et al*. study (Supplementary file 6). Orthologous regions for the *PTCHD3* DMR were also detected to be partially methylated in all three mammals but the *FANCC* DMR was only partially methylated in chimp. These gDMRs could potentially explain the missing ICR for ASE of these genes. The gDMR at the *PTCHD3* promoter can directly suppress maternal allele. *FANCC* gDMR overlaps to a CpG island and CTCF binding site. CTCF is a methylation sensitive DNA-binding protein and CpG methylation can inhibit CTCF binding (Hashimoto et al., 2017; Renda et al., 2007). Moreover, CTCF binding to the first intron of major immediate-early gene of the human cytomegalovirus (HCMV) in HCMV-infected cells resulted in repression of this gene (Puerta et al., 2014). Therefore, the maternally methylated DMR in intron 1 of maternally expressed *FANCC* suggests a mechanism through which paternal allele is suppressed by CTCF binding at DMR while DNA methylation inhibits CTCF binding at maternal allele.

Overall, our study represents a near-complete genome-wide map of human allele-specific methylation by leveraging long-read nanopore technology. This allowed us to expand the set of reported imprinted DMRs using just 12 LCLs with parental SNPs and explain novel DMRs as potential ICRs for several imprinted genes with unknown ICR. 43% of the novel DMRs demonstrated partial methylation in other mammals suggesting their conservation. We detected seven large PofO bias methylation blocks spanning multiple kilobasesd and displaying several features in common. We have suggested two avenues of further investigation: 1) Looking for tissue and individual polymorphism in imprinting, and 2) determining the mechanism and function of the subtle parental bias blocks. We have also shown that nanopore sequencing is a cheap and easy way to call ICRs and can open the way to answering those questions in future. This study provides a blueprint for further surveys using nanopore sequence data and demonstrates the potential of this approach to study personalized allelic methylation in disease such as cancer with wide spread allelic methylation aberrations.

## Materials and Methods

### Nanopore Sequencing Data and Detection of Allele-Specific Methylation

We used publicly available nanopore sequencing data for 12 LCLs with trio data available. Raw and basecalled nanopore data for HG002, HG005, HG00733, HG01109, HG01243, HG02055, HG02080, HG02723, HG03098, and HG03492 obtained from Human Pangenomics and GIAB (Shafin et al., 2020; Zook et al., 2016). NA19240 data (ERR3046934 and ERR3046935 raw nanopore and their basecalled reads ERR3219853 and ERR3219854) obtained from De Coster et al (De Coster et al., 2019). Raw and basecalled nanopore data for NA12878 obtained from rel6 nanopore WGS consortium (Jain et al., 2018). Basecalled reads mapped to GRCh38 using Minimap2 with the setting *minimap2 –ax map-ont* (Kent et al., 2002; Li, 2018). Subsequently, CpG methylations were called using nanopolish with default parameters (Simpson et al., 2017). Methylation calls for each sample preprocessed using NanoMethPhase *methyl_call_processor* default setting for downstream analysis (Akbari et al., 2021). To detect allelic methylation we used high quality SNVs for each cell line and it’s parents. For all the cell lines and parents, except HG002 and HG005, high quality SNVs were called using Strelka2 with default parameters from alignment files in the 1KGP GRCh38 (Auton et al., 2015; Kim et al., 2018). High quality SNVs for HG002 and HG005 and their parents were obtained from GIAB v.3.3.2 high confidence variant calls (Zook et al., 2019). For each LCL a mock phased vcf file with defined parent of origin of each high-quality heterozygous SNV was created using an in-house bash script (https://github.com/vahidAK/NanoMethPhase/tree/master/scripts: Trio_To_PhaseVCF_4FemaleChild.sh & Trio_To_PhaseVCF_4MaleChild.sh). Subsequently, we detected haplotype methylome in each sample using NanoMethPhase with the setting *nanomethphase phase –mbq 0*. Finally, DMRs between haplotypes were called using default setting of NanoMethPhase *dma* module that uses Dispersion Shrinkage for Sequencing data (DSS) R package for DMA and performs a Wald test at each CpG site (Park and Wu, 2016). To avoid the confounding effects of X-inactivation, and because previous studies demonstrated no evidence of imprinting at sex chromosomes, we only examined autosomal chromosomes (Court et al., 2014; Joshi et al., 2016; Zink et al., 2018).

### WGBS Data and Detection of Novel DMRs

To validate allelic methylation in other tissues and also detect potential novel ICRs we used 24 public WGBS (GSM1010978, GSM1010979, GSM1010980, GSM1010981, GSM1010983, GSM1010984, GSM1010986, GSM1010987, GSM1010988, GSM1010989, GSM1112838, GSM1120321, GSM1120326, GSM1127054, GSM1127125, GSM916049, GSM916050, GSM983645, GSM983647, GSM983648, GSM983649, GSM983650, GSM983651, GSM983652) for 20 tissue type samples from Epigenomics Roadmap including adipose, adrenal gland, liver, aorta, brain hippocampus, breast luminal epithelial, breast myoepithelial, esophagus, gastric, left ventricle, lung, ovary, pancreas, psoas muscle, right atrium, right ventricle, sigmoid colon, small intestine, spleen, and thymus (Bernstein et al., 2010). Wig files which include fractional methylation data were obtained and converted to bed format using UCSC tools and lifted over to hg38 coordinates using CrossMap and UCSC lift over chain file (Kent et al., 2002; Zhao et al., 2014). All, bed format files were then merged to keep CpGs that are common in at least 10 samples. At imprinting control regions only one allele is methylated and we expect to observe partial methylation (∼50%) at such regions. However, the adjacent sites which are not imprinted display ∼0% or ∼100% methylation. Therefore, we used a comparison between detected DMRs with their adjacent sites in WGBS data. For each DMR we determined the number of CpG sites with methylation rates between 30-70% (partial methylation) and normalized it by dividing the numbers to all CpGs in the interval. We also determined this ratio for the adjacent sites (>=20kb away and not been reported as imprinted gene or ICR). We then used limma’s linear model to perform statistical analysis of the ratios at each DMR and adjacent sites (Codes are available on https://github.com/vahidAK/NanoMethPhase/tree/master/scripts: PartialCpGMethylationAtDMRandAdjacent.py & ComparePartialMethylationAtDMRsToAdjacentUs ingLimma.R). As controls and because ICRs are usually overlapped with CpG islands, we examined 200 randomly selected CpG islands and 200 randomly selected 1kb, 2kb, and 3kb intervals with more than 15 CpGs.

### Detection of Germline and Somatic DMRs

If a DMR is germline, it is established during germ cell development and survived the pre-implantation methylation reprograming. Therefore, gDMRs will overlap with DMR detected from oocyte vs sperm with consistent methylation direction, i.e. maternally methylated DMRs display high methylation in oocyte and very low or no methylation in sperm and vice versa. Moreover, gDMRs need to display partial methylation after fertilization and early development.

In order to discriminate gDMRs from somatic, we used public WGBS data for 3 sperm and 2 oocyte, and 1 blastocysts libraries and 3 fetal tissue types (GSM1172595 thymus, GSM1172596 muscle, GSM941747 brain) (Bernstein et al., 2010; Okae et al., 2014). Read counts for methylated and unmethylated CpG sites were obtained for sperm and oocyte samples and DMA was performed using NanoMethPhase *dma* module. To detect potential candidate gDMRs, we overlapped detected DMRs from oocyte vs sperm DMA to detected imprinted DMRs from nanopore. We further used blastocysts and fetal tissues to investigate if potential gDMRs escaped the second round of methylation reprograming and if sDMRs are stablished during somatic development.

### Allelic H3K4me3 Analysis

H3K4me3 ChIP-seq fastq files were obtained for NA12878, NA12891, NA12892, NA19238, NA19239, NA19240, and NA18507 (SRA: SRP044271) and aligned to the GRCh38 reference genome using bwamem default setting (Adoue et al., 2014; Kent et al., 2002; Li and Durbin, 2009). High quality SNVs were called for these samples from 1KGP GRCh38 alignment files using strelka2 (Auton et al., 2015; Kim et al., 2018). We then counted the number of reads with minimum mapping quality of 20 and base quality of 10 at each heterozygous SNV and kept those with more than 5 mapped reads for binomial test. The reference allelic counts and total counts at each heterozygous SNV (or maternal allelic counts and total counts in case for trios) were used to detect significant allelic bias using a two-sided binomial test under the default probability of *P = 0*.*5* in python SciPy package (Codes are available on https://github.com/vahidAK/NanoMethPhase/tree/master/scripts: CountReadsAtSNV.py & Binomial_test.py) (Virtanen et al., 2020).

### Mammalian Conservation of DMRs

To test whether any of the detected novel DMRs are conserved in other mammals we used 17 WGBS datasets for mus musculus (GSE42836, includes 17 tissue types), 34 WGBS datasets for rhesus macaque (GSE34128, includes 6 peripheral blood samples. GSE151768 includes 15 NeuN+ and 13 OLIG2+ brain samples), and 22 WGBS datasets for chimpanzee (GSE151768, includes 11 NeuN+ and 11 OLIG2+ brain samples) to examine partial methylation in orthologous intervals (Hon et al., 2013; Jeong et al., 2021; Tung et al., 2012). Mouse, Macaque, and Chimp coordinates lifted over to mm10, RheMac8, and PanTro5 coordinates using CrossMap and appropriate liftover file from UCSC, if they were not already in this coordinates. The list of detected human DMRs were also converted to the orthologous regions for each mammal using CrossMap and the appropriate UCSC liftover file (Kent et al., 2002; Zhao et al., 2014). Since many coordinates in human splinted to several orthologous in other mammals, we merged orthologous intervals which were <=200bp apart. Finally, we used our approach explained in aforementioned section (WGBS Data and Detection of Novel DMRs) to detect orthologs with significant partial methylation.

To examine the somatic and germline orthologous DMRs, we used two embryo (GSM3752614, GSM4558210), one sperm (GSE79226, combined methylation frequencies from three replicates), and three oocyte (GSM3681773, GSM3681774, GSM3681775) WGBS libraries from mouse; And one embryo (GSM1466814), one sperm (GSM1466810), and one oocyte (GSM1466811) WGBS libraries from rhesus macaque (Dahlet et al., 2020; Gao et al., 2017; Jung et al., 2017; Saenz-de-Juano et al., 2019).

## Acknowledgements

SJMJ and MAM acknowledge funding from the Canada Research Chairs program and the Canadian Foundation for Innovation. VA acknowledges funding from the University of British Columbia Four Year Doctoral Fellowship.

## Supplementary Files

**Supplementary File 1:** List of the 383 reported imprinted DMRs from previous studies.

**Supplementary File 2:** Detected DMRs using nanopore sequencing data across 12 LCLs and their overlap to reported DMRs.

**Supplementary File 3:** Examining significance of partial methylation of the detected DMRs from nanopore in 24 WGBS datasets from 20 tissue types. Examining additional 200 randomly selected CGIs, 1kb, 2kb, and 3kb intervals as controls. DNA methylation of the DMRs and their up - and down-stream regions at each WGBS sample.

**Supplementary File 4:** Investigating the significance of partial methylation of the undetected reported DMRs in 24 WGBS datasets from 20 tissue types.

**Supplementary File 5:** Mapping of the undetected reported DMRs to DMRs detected in each nanopore LCL sample.

**Supplementary File 6:** List of the DMRs detected from nanopore data and validated using WGBS data. Average methylation at 107 nanopore-detected DMRs and validated with WGBS data in LCLs, fetal, embryos and gamete samples.

**Supplementary File 7:** Allelic counts for each heterozygous SNV at the detected DMRs and binomial test results of H3K4me3 ChIP-seq in 7 LCL samples. Allelic counts for each PofO defined SNV at the detected DMRs and binomial test results of H3K4me3 ChIP-seq in NA12878 and NA19240 samples.

**Supplementary File 8:** Limma results for partial methylation of human DMRs’ orthologs in chimpanzee, rhesus macaque, and mus musculus WGBS data. Average DNA methylation of orthologs in mouse and rhesus macaque adult tissues and their embryos and gametes.

**Supplementary File 9:** List of the 259 known imprinted genes.

**Supplementary File 10:** Mapping detected DMRs to close vicinity of the known imprinted genes.

## Supplementary Figures

DMRs covered by each nanopore LCL sample and various IGV screenshots of the novel imprinted DMRs and blocks.

