## Supplementary Figures for "Genome-Wide Detection of Imprinted Differentially Methylated Regions Using Nanopore Sequencing"

**Note 1**: In all IGV screenshots, PofO ASE track represents allele-specific expression track from Zink *et al.*, (https://www.nature.com/articles/s41588-018-0232-7) without any filtering for *P value*, which is:

(log(#reads with ref allele / #reads with alt allele) when paternal homologue has ref allele - log(#reads with ref allele / #reads with alt allele) when paternal homologue has alt allele)/2

Positive or upward bars represent paternal expression bias and negative or downward bars represent maternal expression bias.

**Note 2**: In all IGV screenshots, Adult Tissues track represents the average methylation of 24 WGBS data from 20 tissue samples and Fetal Tissues track is the average methylation of 3 fetal WGBS tissue samples. These WGBS samples obtained from Roadmap Epigenomics Project (https://www.ncbi.nlm.nih.gov/geo/roadmap/epigenomics/).

**Note 3**: In all IGV screenshots, blastocyst, oocyte and sperm tracks are the methylation from one blastocysts, and average methylation from two oocyte, and three sperm libraries from Okae et al., 2014. (https://doi.org/10.1371/journal.pgen.1004868)

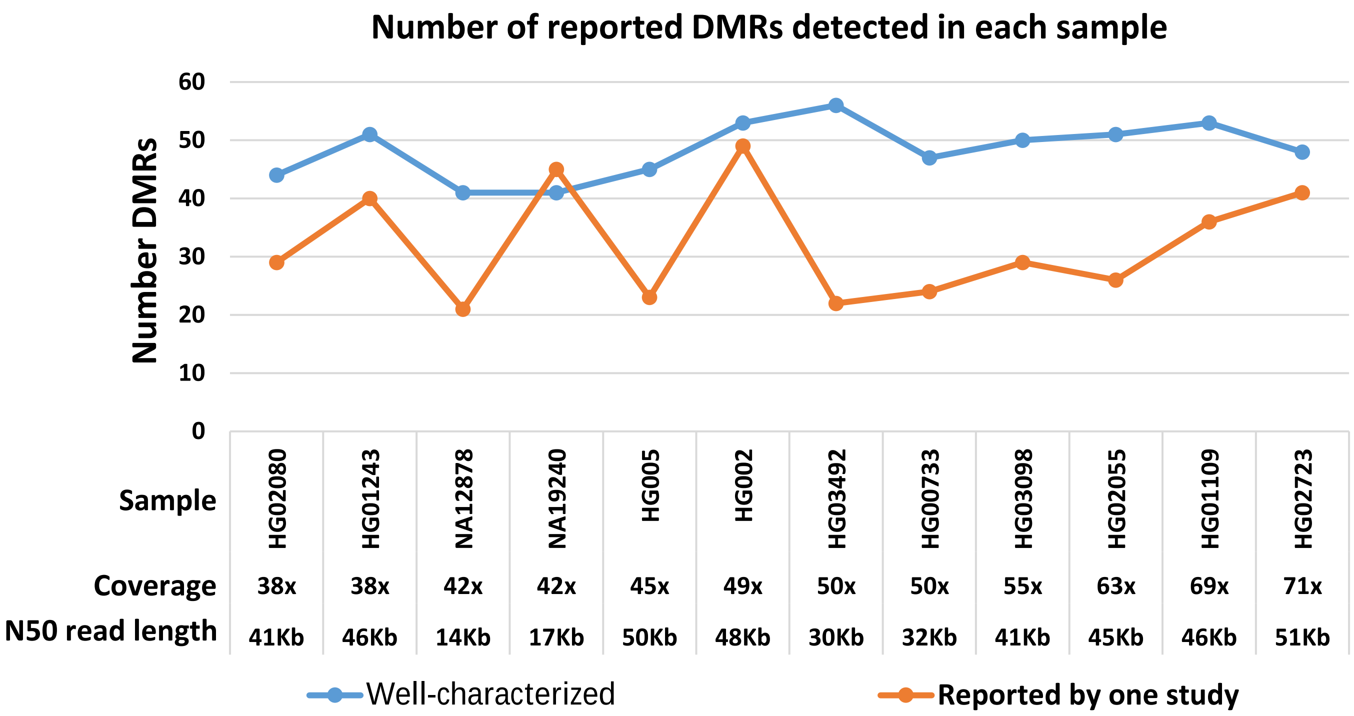


Figure S1: Number of allelic DMRs in each sample overlapped to the reported DMRs.

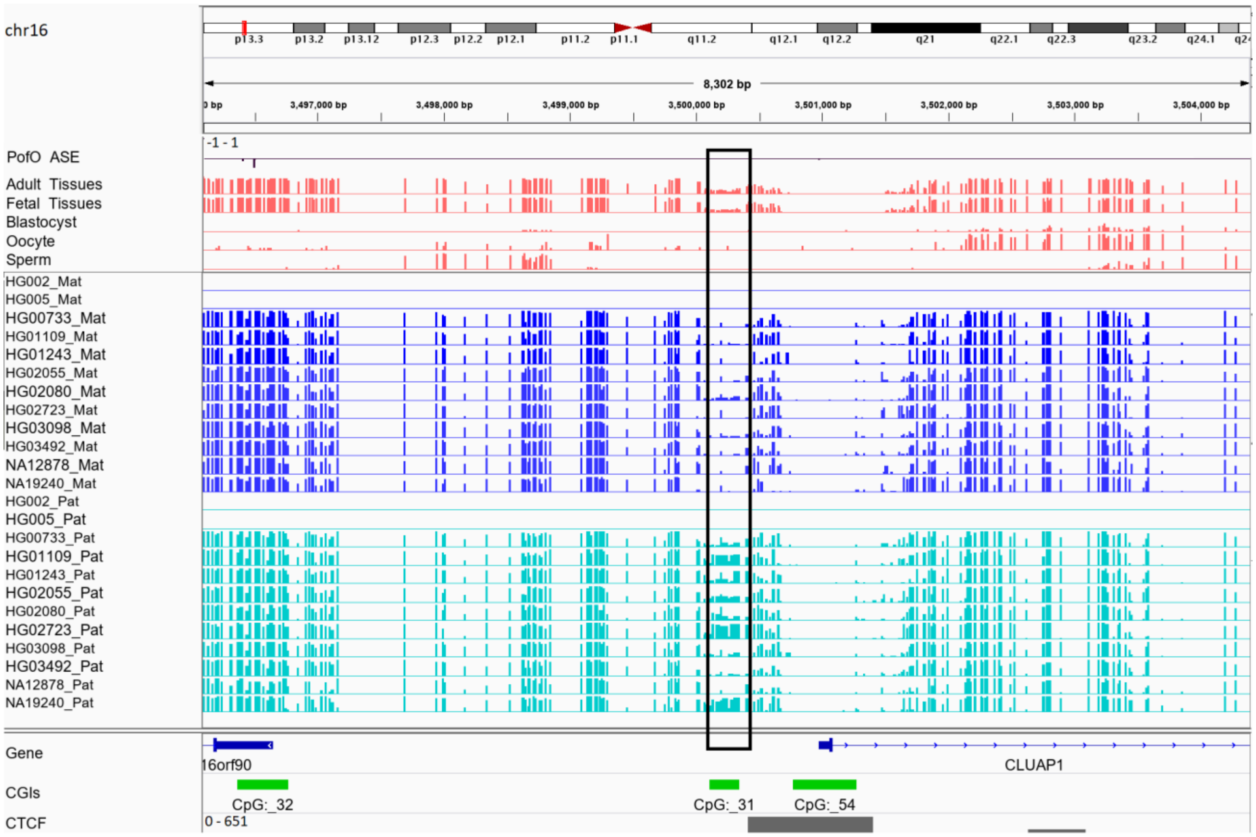


Figure S2: Paternally methylated novel somatic DMR ~13 Kb downstream of the maternally expressed *NAA60* gene. DMR is in the black box. Range for all methylation tracks is from 0-1.

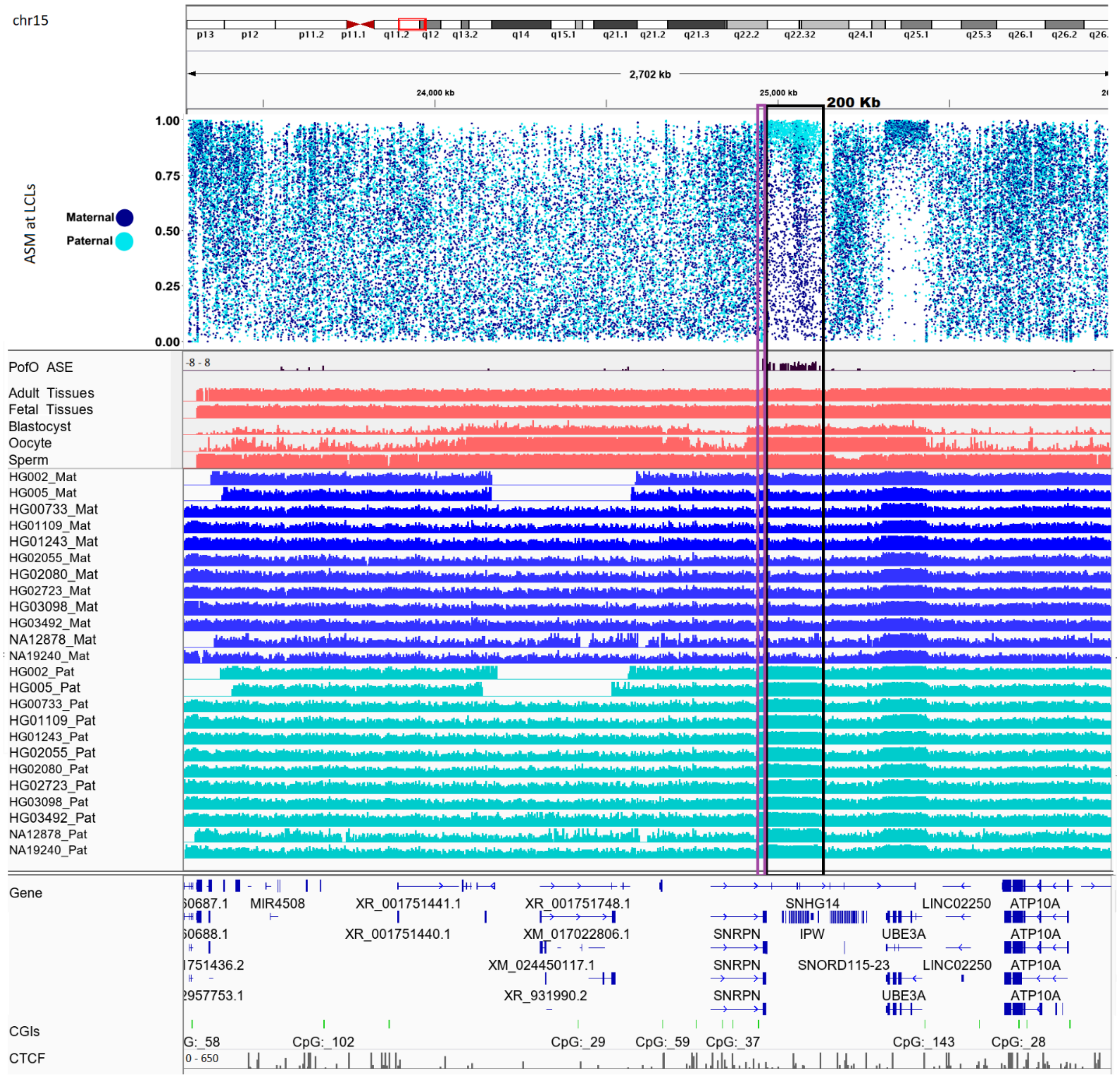


Figure S3: Continuous 200Kb paternally methylated and expressed block at *PWS/AS* cluster. Black box represents whole block and purple box represents germline DMR. Range for all methylation tracks is from 0-1. To better demonstrate subtle allelic methylation bias, we also represented methylation from LCLs as point plot.

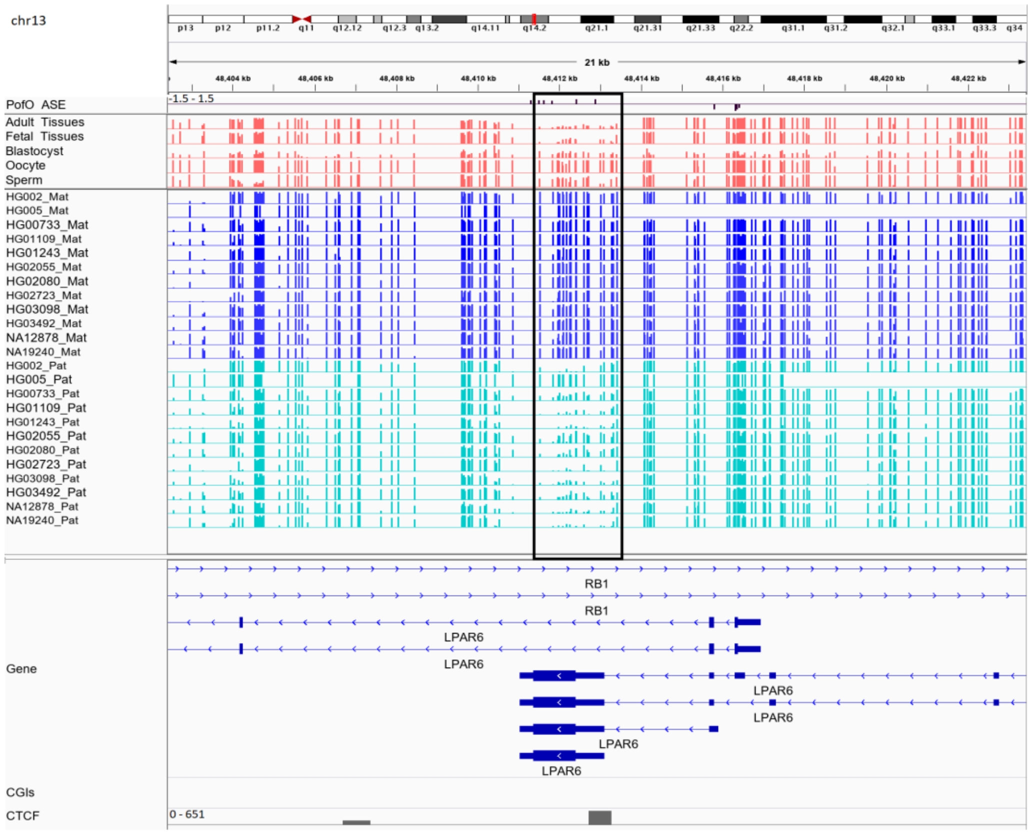


Figure S4: Novel somatic maternally methylated DMR in maternally expressed RB1 gene and Isoform dependent imprinted LPAR6 gene. DMR is in the black box. Range for all methylation tracks is from 0-1.

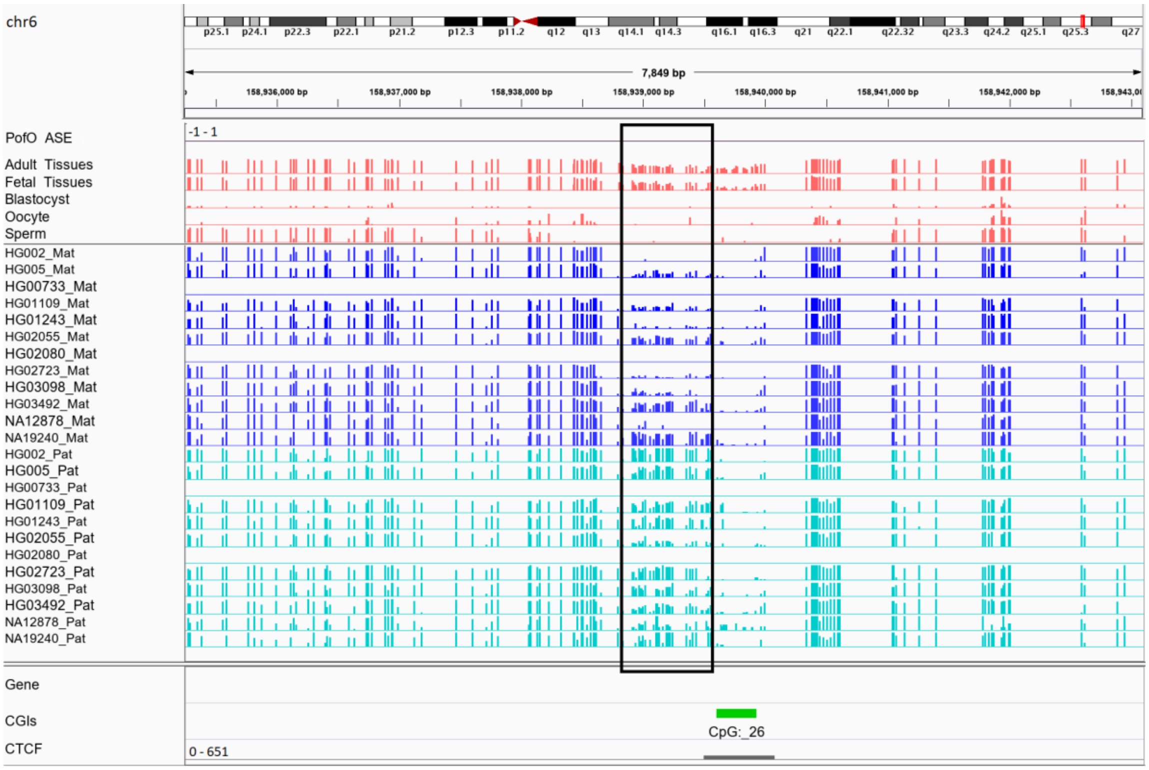


Figure S5: Novel somatic paternally methylated DMR ~1 Mb upstream maternally expressed IGF2R gene. DMR is in the black box. Range for all methylation tracks is from 0-1.

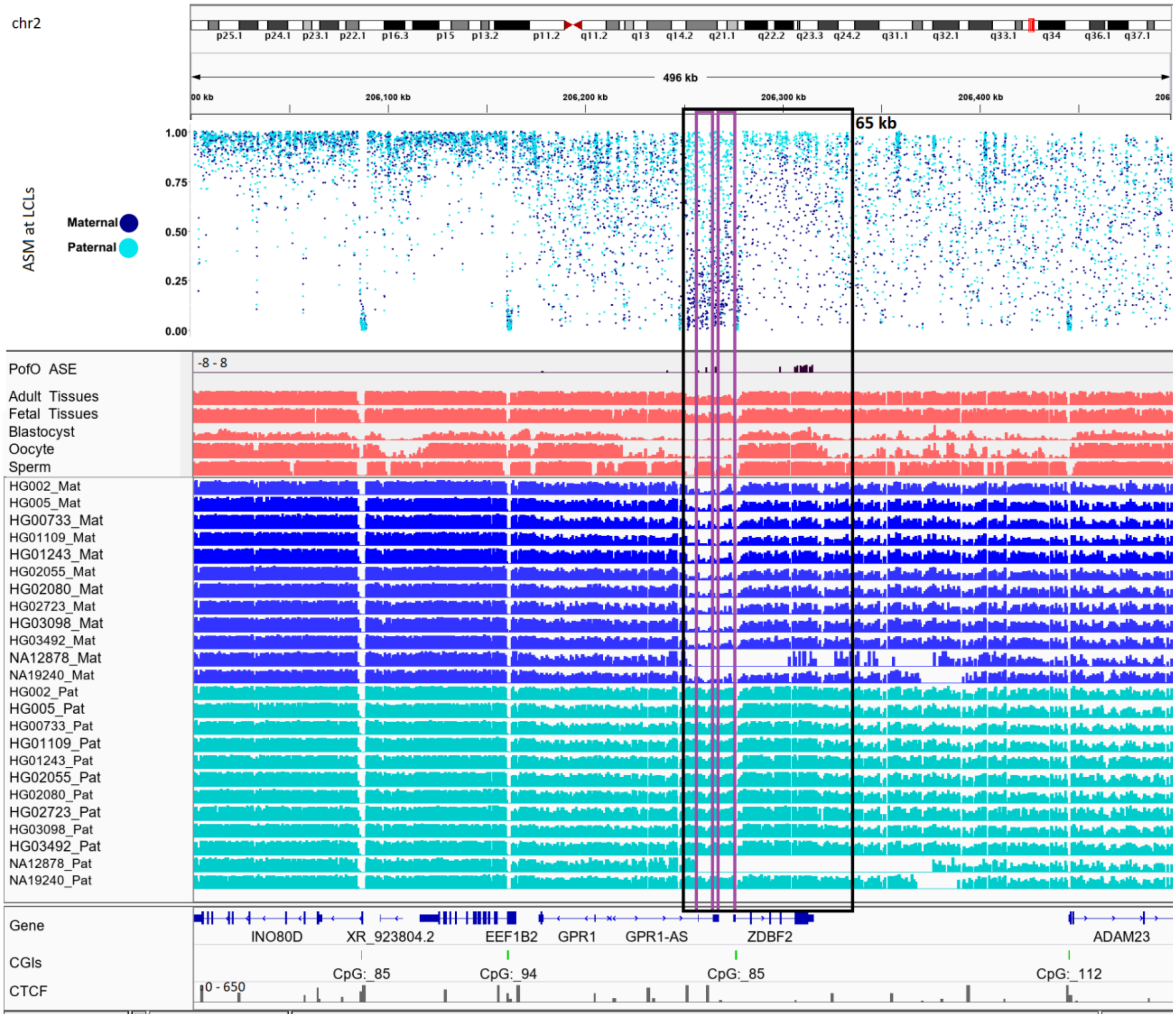


Figure S6: Continuous 65Kb paternal methylation bias and paternal expressed block at *GPR1-AS/ZDBF2* cluster. Black box represents whole block and purple box represents germline DMR. Range for all methylation tracks is from 0-1. To better demonstrate subtle allelic methylation bias, we also represented methylation from LCLs as point plot.

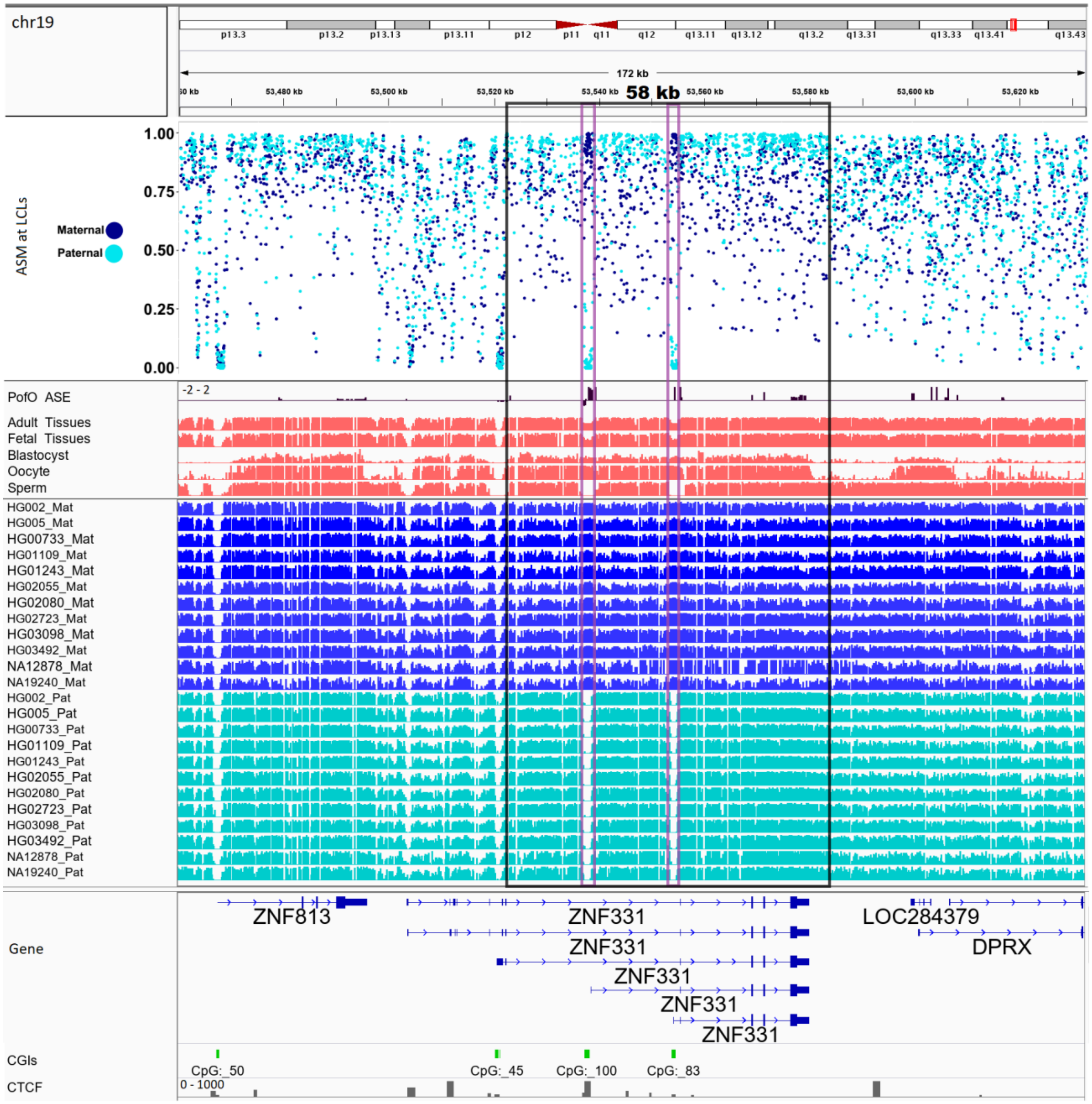


Figure S7: Continuous 58Kb parental methylation bias and paternal expressed block at *ZNF331/ZNF813* cluster. Black box represents whole block and purple box represents germline DMR. Range for all methylation tracks is from 0-1. To better demonstrate subtle allelic methylation bias, we also represented methylation from LCLs as point plot.

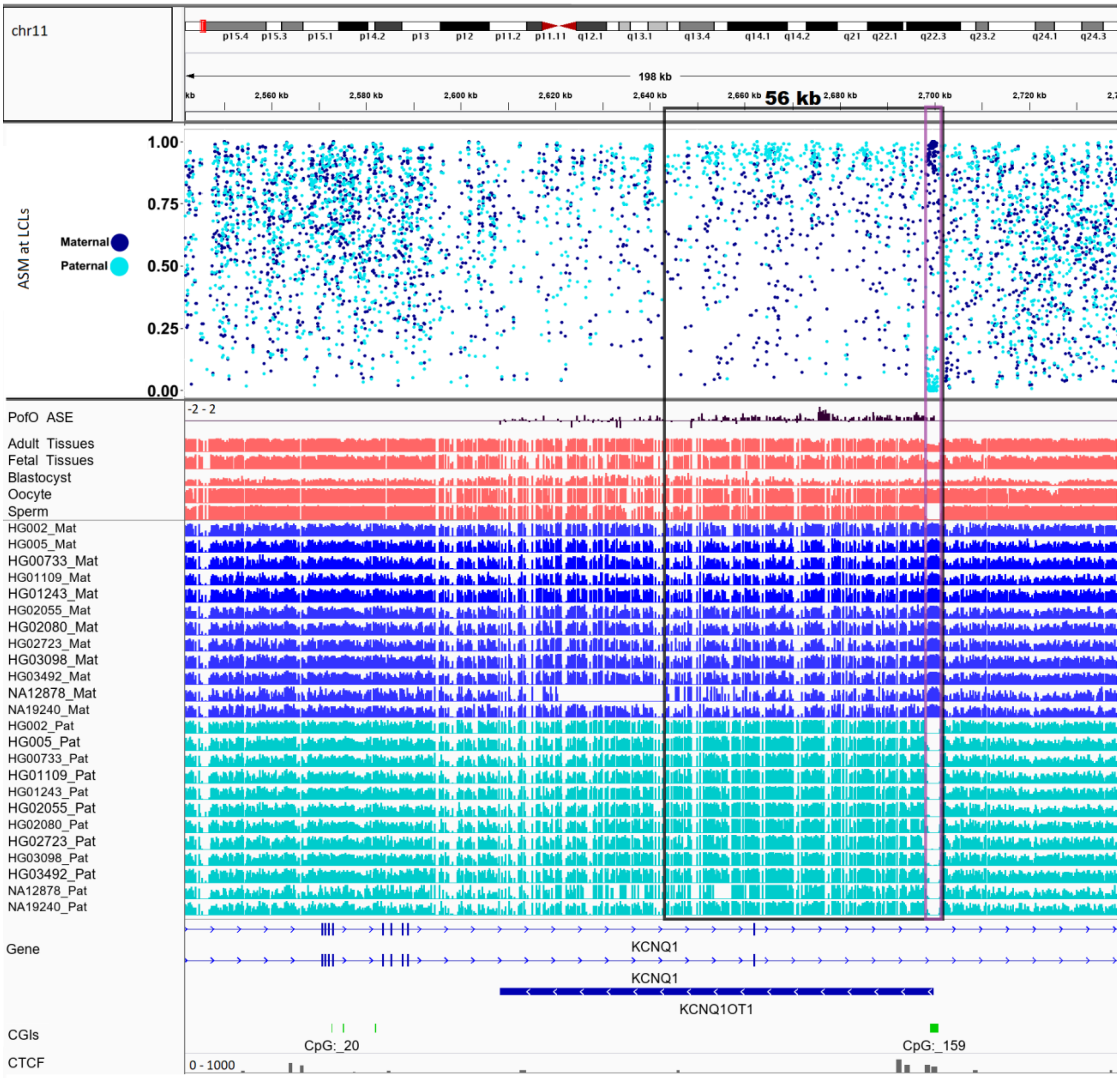


Figure S8: Continuous 56Kb parental methylation bias and paternal expressed block at *KCNQ1/KCNQ1OT1* cluster. Black box represents whole block and purple box represents germline DMR. Range for all methylation tracks is from 0-1. To better demonstrate subtle allelic methylation bias, we also represented methylation from LCLs as point plot.

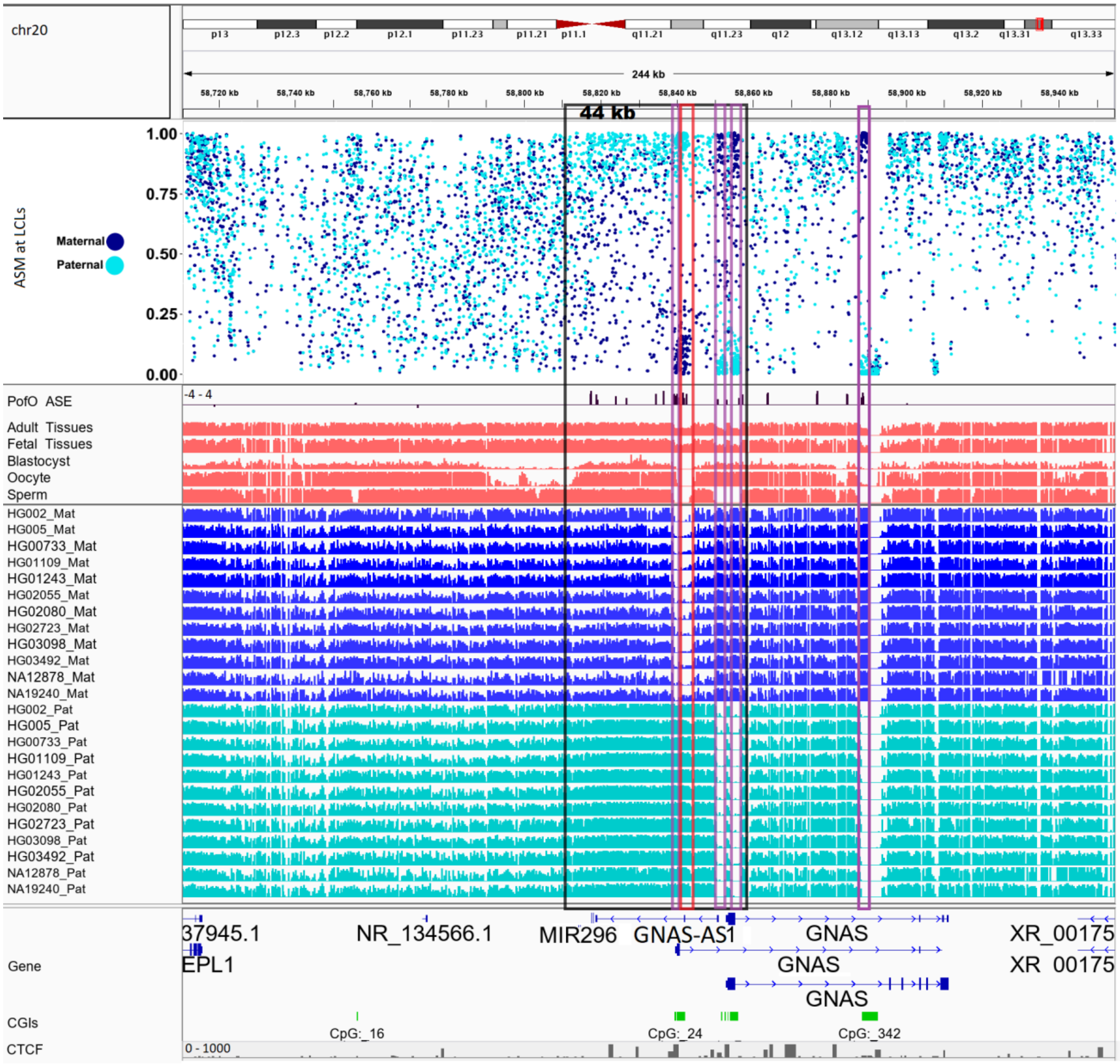


Figure S9: Continuous 44Kb parental methylation bias and paternal expressed block at *GNAS* cluster. Black box represents whole block and purple box represents germline DMR and red box represents somatic DMR. Range for all methylation tracks is from 0-1. To better demonstrate subtle allelic methylation bias, we also represented methylation from LCLs as point plot.

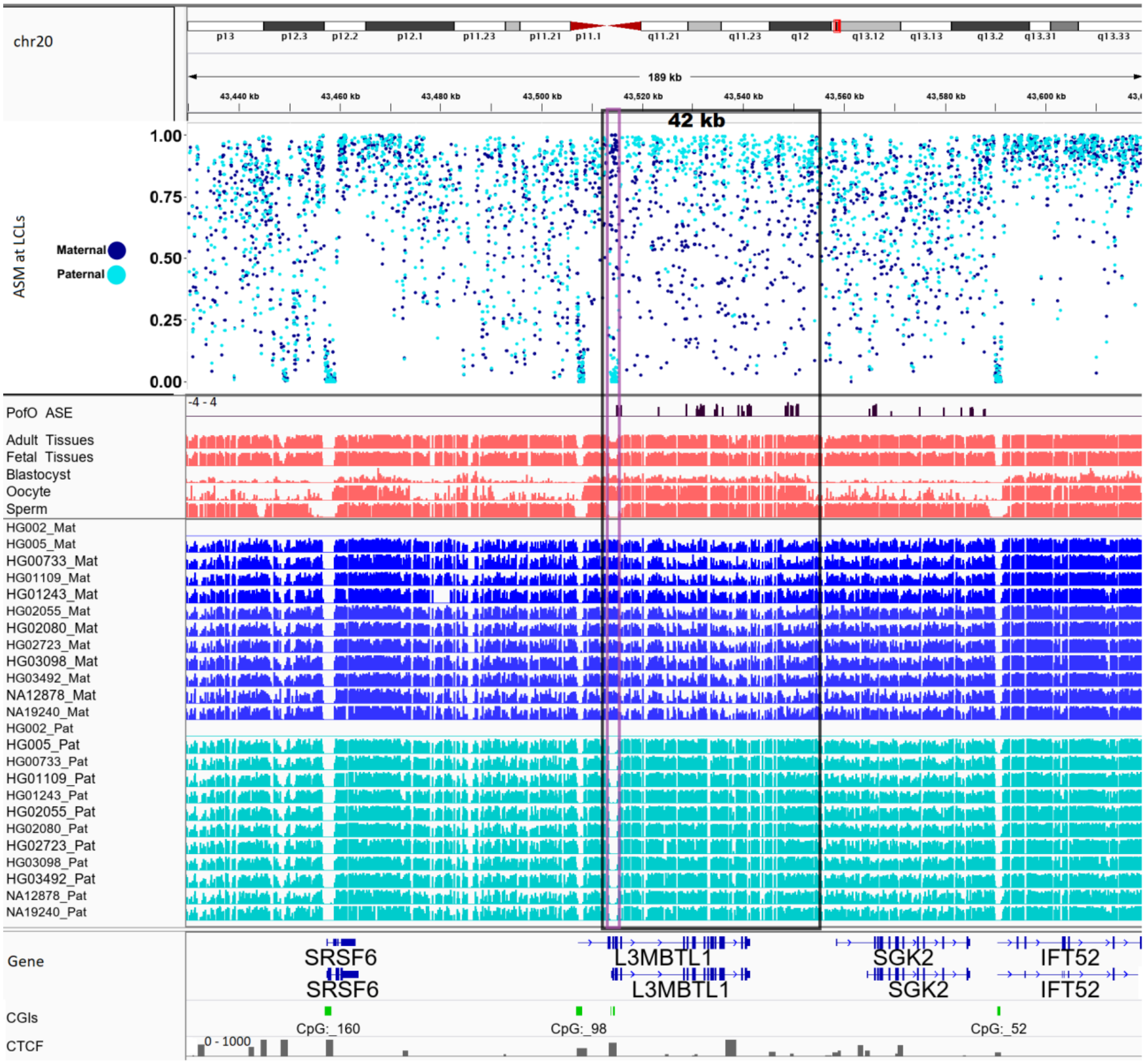


Figure S10: Continuous 42Kb parental methylation bias and paternal expressed block at *L3MBTL1/SGK2* cluster. Black box represents whole block and purple box represents germline DMR. Range for all methylation tracks is from 0-1. To better demonstrate subtle allelic methylation bias, we also represented methylation from LCLs as point plot.

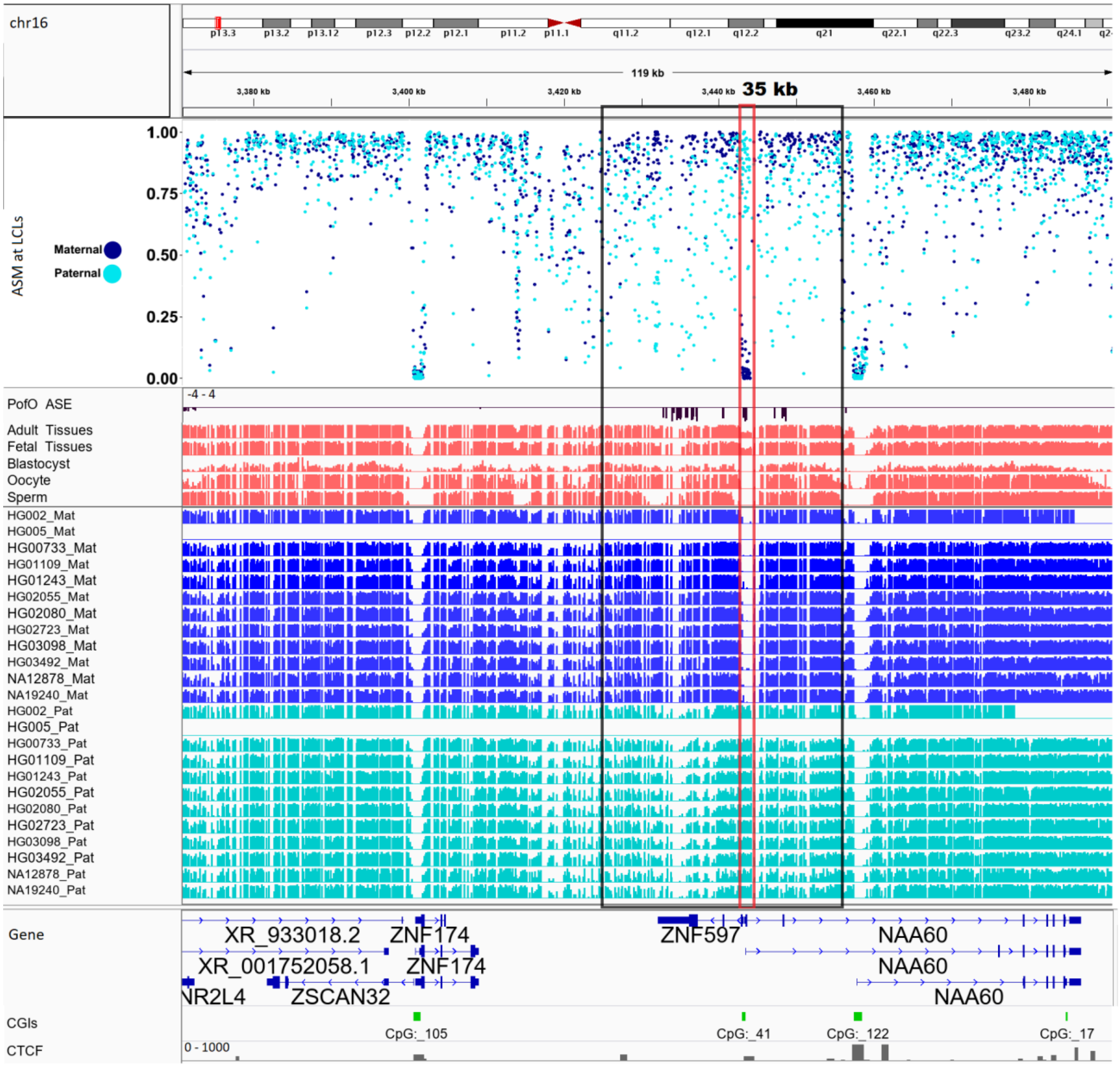


Figure S11: Continuous 35 Kb parental methylation bias and maternal expressed block at *ZNF597/NAA60* cluster. Black box represents whole block and red box represents somatic DMR. Range for all methylation tracks is from 0-1. To better demonstrate subtle allelic methylation bias, we also represented methylation from LCLs as point plot.

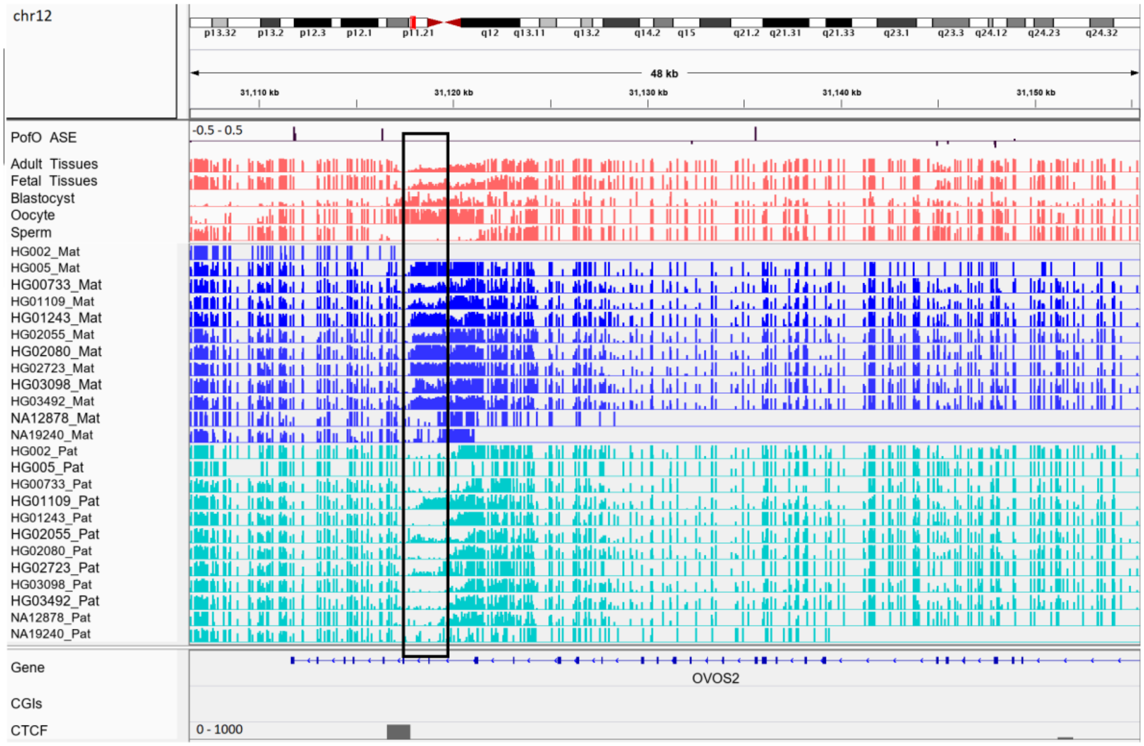


Figure S12: Maternally methylated novel germline DMR in paternally expressed *AC024940.1 (OVOS2)* gene. DMR is in the black box. Range for all methylation tracks is from 0-1.

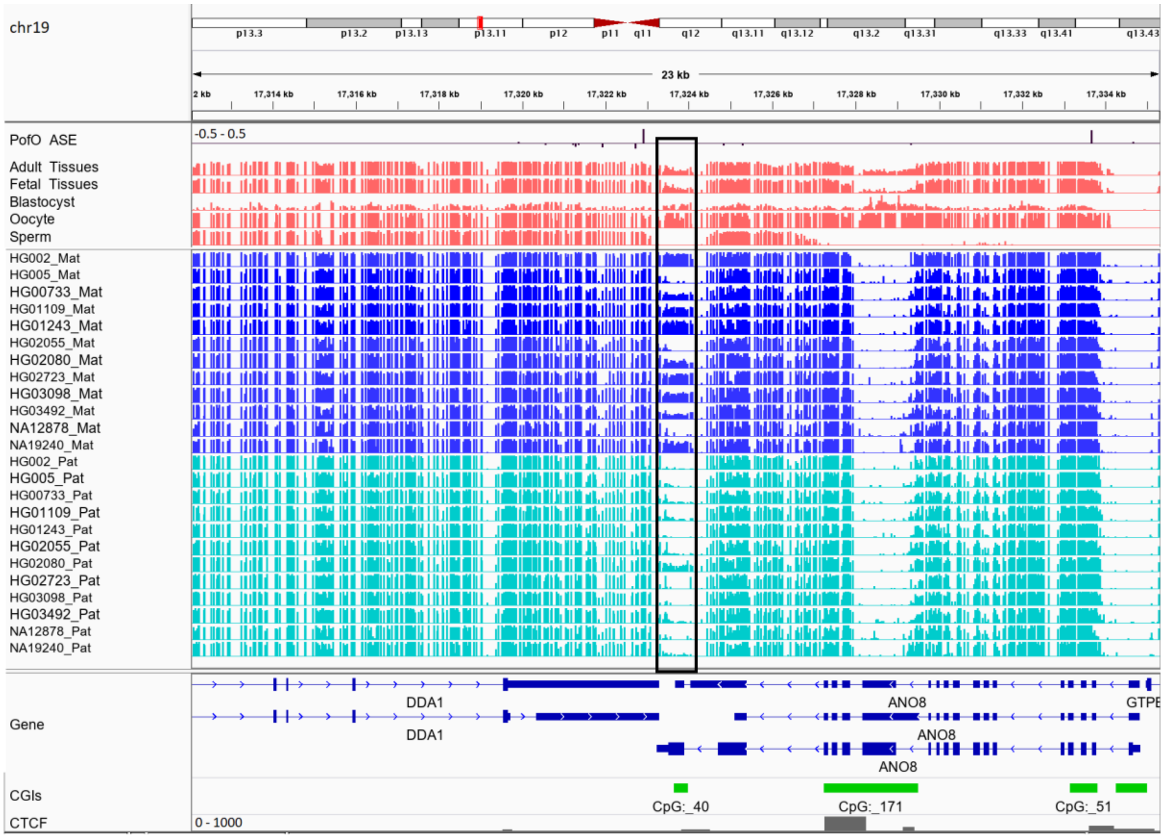


Figure S13: Maternally methylated novel germline DMR at maternally expressed *DDA1* gene. DMR is in the black box. Range for all methylation tracks is from 0-1.

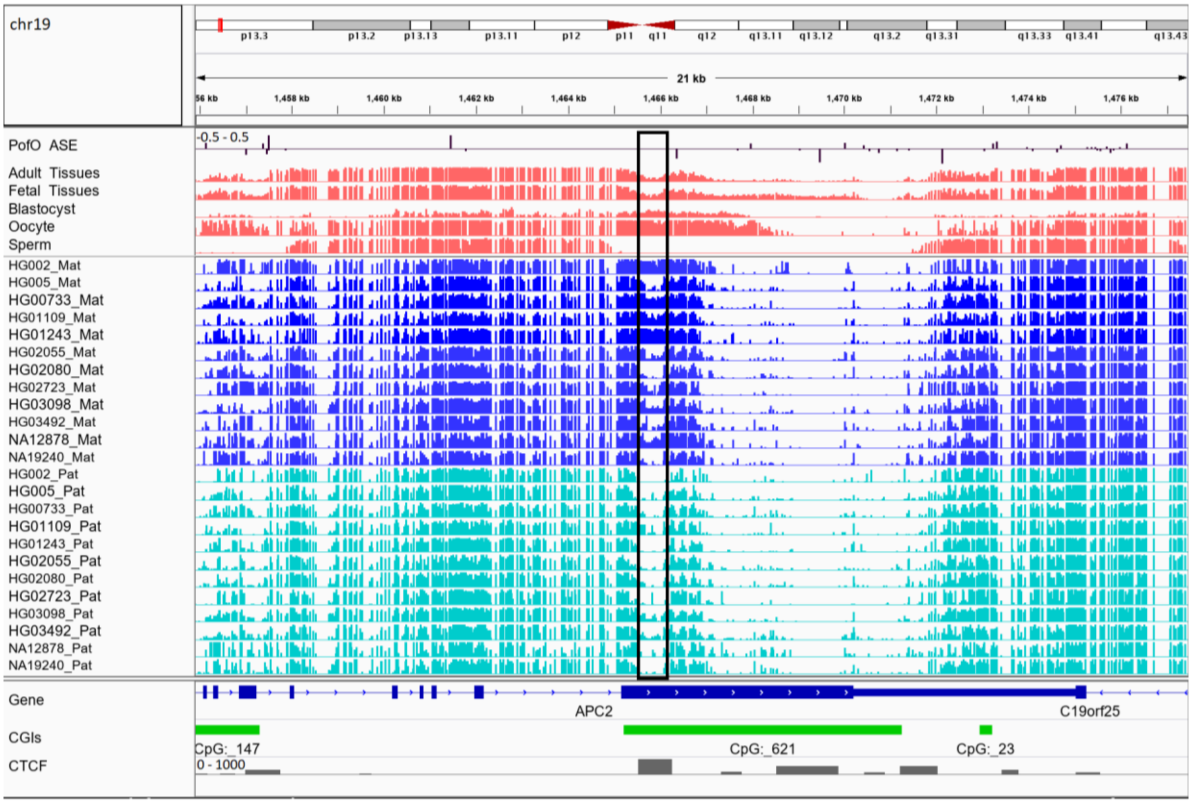


Figure S14: Maternally methylated novel germline DMR ~38 Kb downstream of maternally expressed *ADAMTSL5*. DMR is in the black box. Range for all methylation tracks is from 0-1.

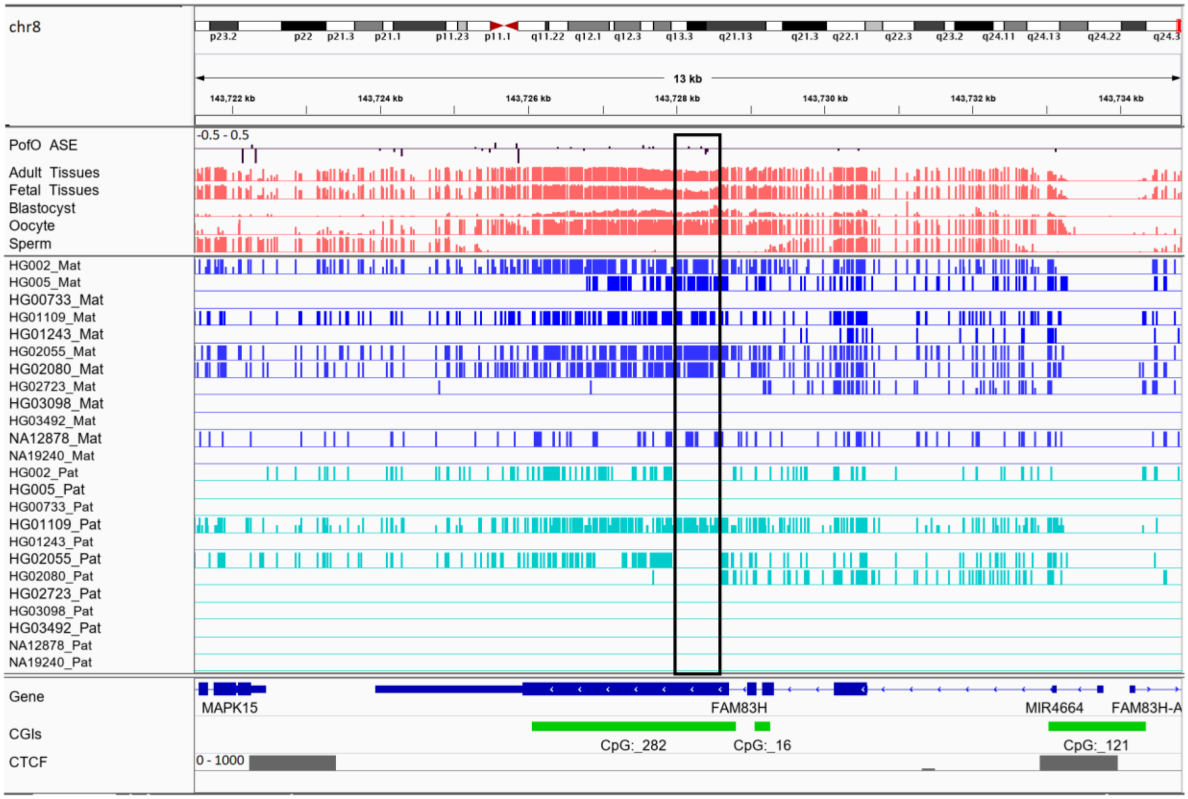


Figure S15: Maternally methylated novel germline DMR ~149 Kb upstream of the isoform dependent imprinted *NAPRT* gene. DMR is in the black box. Range for all methylation tracks is from 0-1.

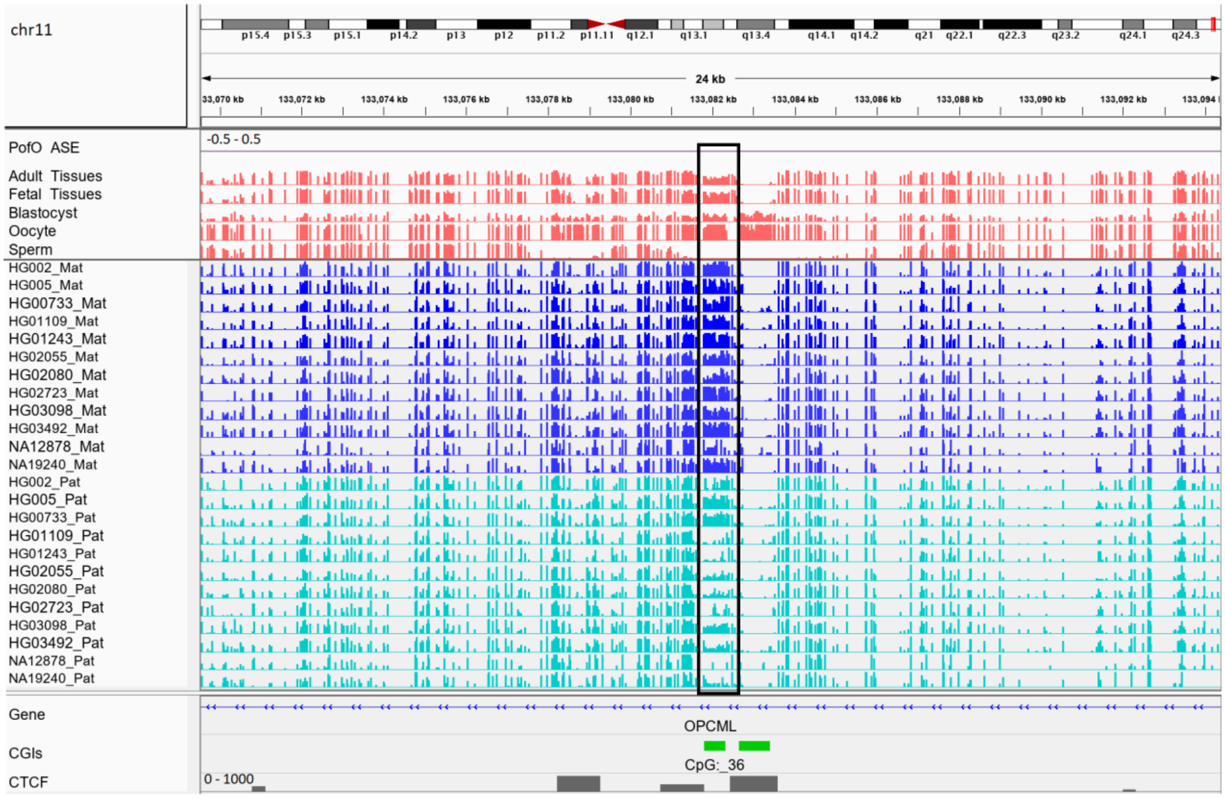


Figure S16: Maternally methylated novel germline DMR ~744 Kb downstream of the maternally expressed *NTM* gene. DMR is in the black box. Range for all methylation tracks is from 0-1.

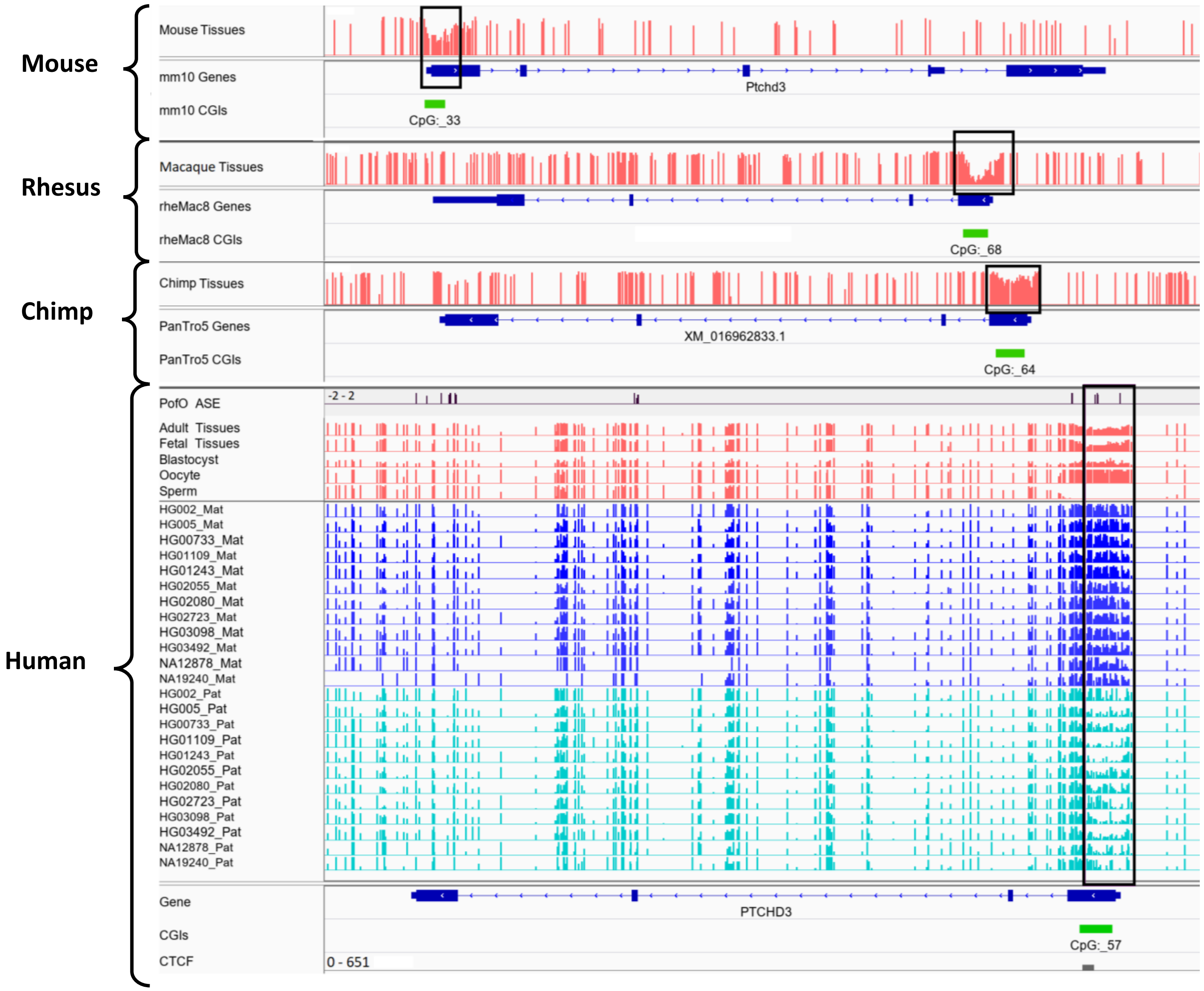


Figure S17: Maternally methylated germline DMR at the promoter of paternally expressed *PTCHD3* gene. The orthologous regions in mouse, rhesus, and chimp are partially methylated which suggest the imprinting at the orthologous regions. Boxes are showing DMRs. Range for all methylation tracks is from 0-1. Mouse, Macaque and Chimp tissues tracks are representing the average methylation from WGBS tissue samples for them.

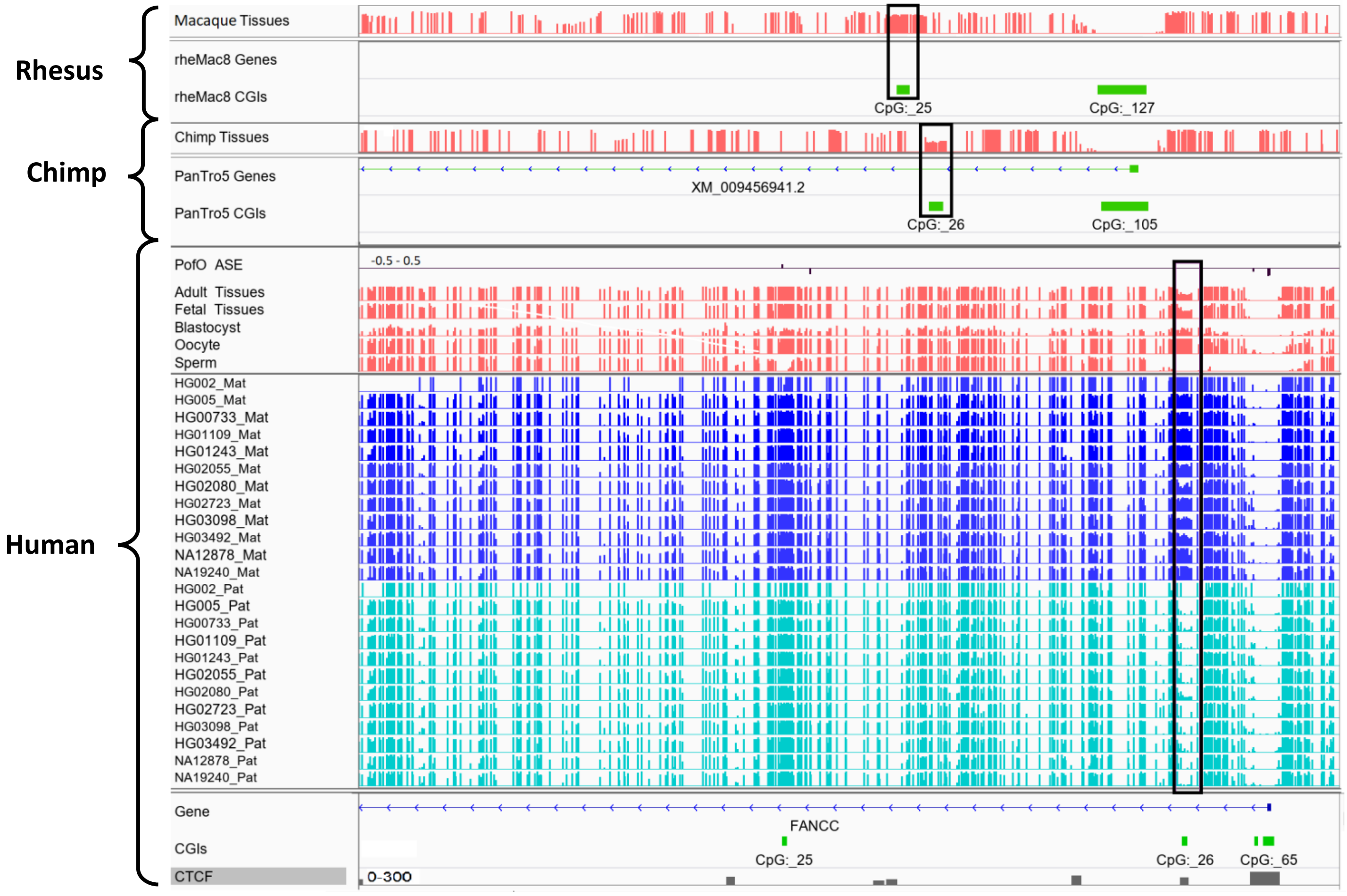


Figure S18: Maternally methylated germline DMR at the intron 1 of maternally expressed *FANCC* gene. The orthologous region in chimp is partially methylated which suggest the imprinting at the orthologous regions and conservation in Chimp. The orthologous region for mouse was not detected. Boxes are showing DMRs. Range for all methylation tracks is from 0-1. Macaque and Chimp tissues tracks are representing the average methylation from WGBS tissue samples for them.

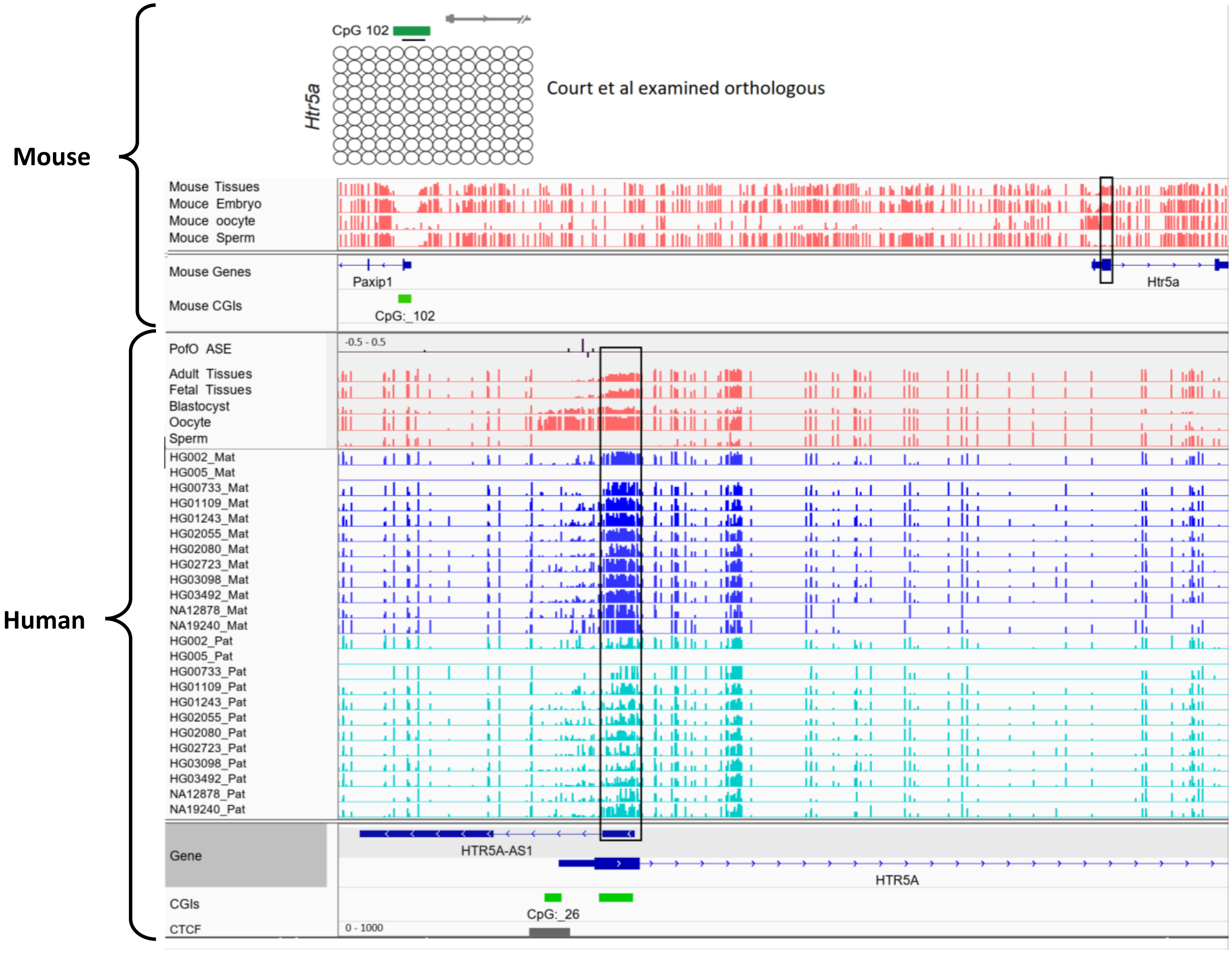


Figure S19: Conserved DMR at *HTR5A* in human and mouse and different orthologous region examined by Court et al. The known DMR at *HTR5A* reported to be not conserved in mouse by court et al (PMID: 24402520), however we detected it as conserved due to different orthologous examined in our study. Court et al examined CGI 102 which is also not imprinted in our analysis, however the orthologue we examined spans beginning of the *HTR5A* and is imprinted with significant partial methylation. Boxes are showing DMRs. Range for all methylation tracks is from 0-1. Mouse tissues, embryo, oocyte and sperm tracks are representing the average methylation from WGBS data.
